# A continuous landscape of signaling encodes a corresponding landscape of CAR T cell phenotype

**DOI:** 10.1101/2025.06.05.658149

**Authors:** Wansang Cho, Jenny Y Liu, Alex N Beckett, Judith C Lunger, Peng Xu, Katie Ho, Ethan E Chen, Antonio Salcido-Alcántar, Lucas E Sant’Anna, Kamal Obbad, Nakoa Po, Elena Sotillo, Crystal L Mackall, Kyle G Daniels

**Affiliations:** Department of Genetics, Stanford University School of Medicine, Stanford, CA 94305, USA; Center for Cancer Cell Therapy, Stanford Cancer Institute, Stanford University School of Medicine, Stanford, CA 94305, USA; Department of Pediatrics, Stanford University School of Medicine, Stanford, CA 94305, USA; Cancer Biology Program, Stanford University, Stanford, CA 94305, USA; Department of Bioengineering, Stanford University, Stanford, CA 94305, USA; Department of Mechanical Engineering, Stanford University, Stanford, CA 94305, USA; Parker Institute for Cancer Immunotherapy, San Francisco, CA 94129, USA; Department of Medicine, Stanford University School of Medicine, Stanford, CA 94305, USA

**Author notes:** These authors contributed equally to this work (Wansang Cho, Jenny Y Liu, Alex N Beckett).

## Abstract

Cytokine signaling is critical to the function of natural immune cells and engineered immune cell therapies such as chimeric antigen receptor (CAR) T cells. It remains unclear how the limited set of signal transducers and activators of transcription (STATs) and other proteins activated by these cytokine receptors can encode the observed diversity of immune cell phenotypes. To understand how signaling downstream of cytokines control immune cell phenotype, we sought to map the structure of Janus kinase (JAK)/STAT signaling domains to cell signaling and resulting CAR T cell function. We recombined 14 signaling motifs to construct a library of ∼30,000 constitutively active synthetic cytokine receptors (SCRs) with intracellular domains composed of novel signaling motif combinations that activate different signaling cascades. We experimentally tested ∼450 SCRs which generated a range of CAR T cell memory, cytotoxicity, and proliferation. SCRs with pSTAT1 and 3 signaling generated effector memory CAR T cells, while SCRs with strong pSTAT5 generated effector CAR T cells with potent anti-tumor activity. To map the structure-signaling-phenotype landscape we trained models to predict signaling and CAR T cell phenotype that result from varied motif combinations. From neural network predictions we identified features, including strong STAT5 and Shc signaling, that promote unsafe autonomous CAR T cell proliferation. Models also revealed a trade-off between memory and cytotoxicity, with a Pareto front encoded by a continuous change in signaling. These results demonstrate that recombination of a limited set of signaling motifs creates a continuous spectrum of signaling that encodes a corresponding spectrum of cell phenotype. This work synthetically expands the combinatorial space of JAK/STAT signaling and provides a foundation for rational design of CAR T cells with improved cytotoxicity, memory, and safety profiles.

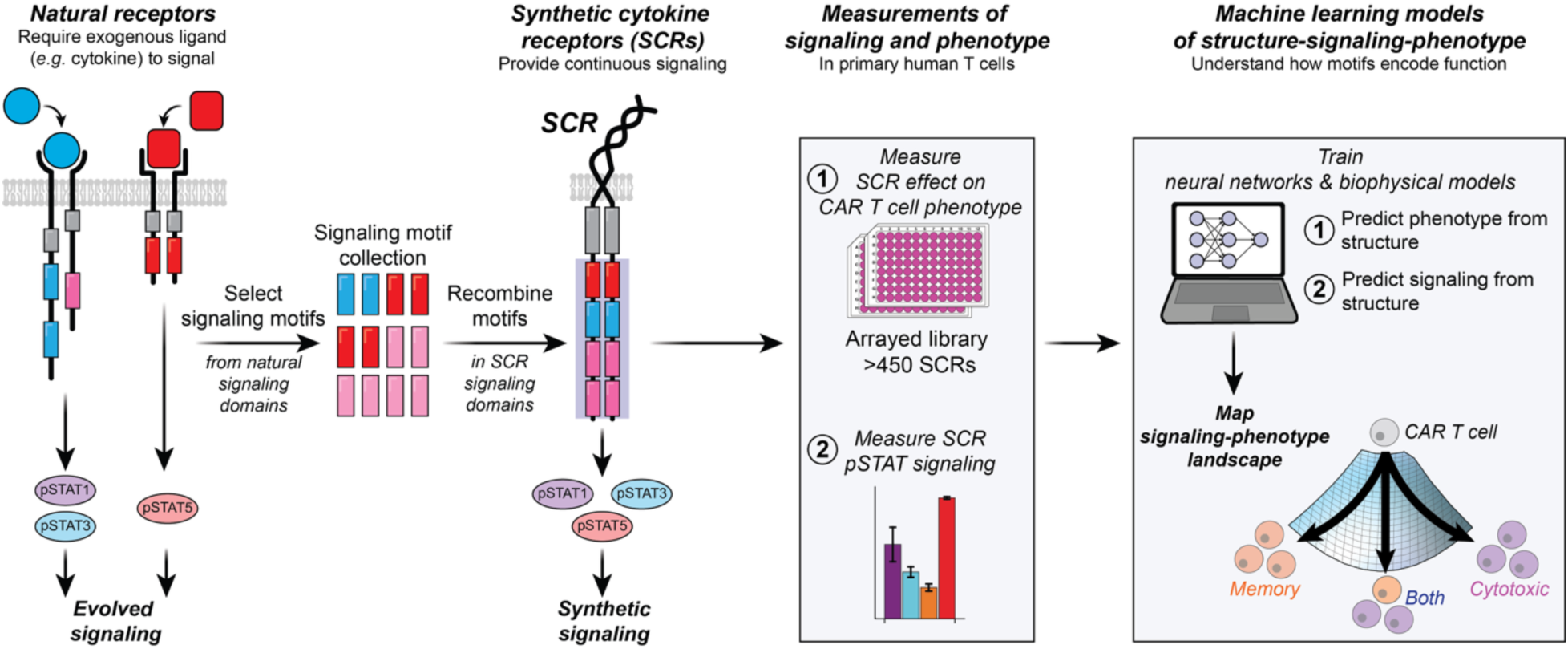

## Introduction

Cytokine-induced JAK/STAT signaling regulates the development, homoeostasis, and function of immune cells(*1, 2*). Aberrant JAK/STAT signaling can cause immune disorders and cancer(*3, 4*). Many studies demonstrate that engineered JAK/STAT signaling has the potential to improve immune cell therapies. More than 50 cytokines and growth hormones have been implicated in JAK/STAT signaling(*5*). These cytokines bind to receptors to activate signaling cascades that orchestrate complex biological processes. Cytokine receptor signaling domains contain signaling motifs that recruit and activate proteins such as STATs, kinases, phosphatases, phospholipases, adapters, and other proteins(*5–8*). It has long been appreciated that the choice of signaling motif determines the STATs and other proteins activated by cytokine receptors(*6*). However, it remains unclear how signaling motifs and the limited set of proteins they activate encode the observed diversity of immune cell phenotypes. Immunology, cancer biology, and immune engineering would benefit from a quantitative understanding that connects signaling domain structure (combinations of conserved motifs), activation of signaling pathways, and resulting immune cell phenotype.

CAR T cells have been effective against an array of blood cancers(*9*), and recent advances have demonstrated the potential of CAR T cells in solid tumors(*10–14*). Considerable effort has been spent on engineering cell signaling, including JAK/STAT signaling to improve CAR T cell efficacy against blood cancers and solid tumors. These efforts include constitutively active STAT (*15–17*), overexpressed cytokine(*18–22*), tethered cytokine(*23, 24*), super agonist cytokine(*25, 26*), orthogonal cytokine signaling(*27, 28*), JAK/STAT CAR signaling domains(*29*), and constitutively active cytokine receptors(*13, 30*). It is understood that the JAK/STAT signaling associated with different cytokines shapes CAR T cell response(*31*). However, a quantitative and predictive understanding of the structure-signaling-phenotype relationship eludes us. For example, which signaling motifs and signaling pathways encode desirable CAR T cell properties like persistence and cytotoxicity, or undesirable properties like autonomous proliferation? Machine learning advances have enabled rational design of structured proteins(*32, 33*), but these advances have not extended to the unstructured signaling domains that mediate JAK/STAT signaling. Similar advances to enable rational design of JAK/STAT signaling domains to tune cell signaling and phenotype would improve cell therapy(*34*). However, these advances will require experimental data that includes a variety of signaling domains, signaling measurements, and cell phenotypes.

Recent work demonstrated the potential of constitutively dimerized cytokine receptors (ZipRs) with natural signaling domains to improve CAR T cell proliferation and anti-tumor activity(*30*). These ZipRs contained signaling domains from interleukin (IL) 2 Receptor (R), IL7R, IL12R, and IL21R. Other recent work showed that recombination of signaling motifs in CAR signaling domains can generate synthetic signaling domains that result in a diverse range of CAR T cell differentiation, proliferation, and cytotoxicity(*35*). We reasoned that constitutively dimerized chimeric cytokine receptors would serve as a powerful platform to create synthetic signaling domains with motif combinations that expand our control of cell function. Here, we created a library of constitutively dimerized cytokine receptors with synthetic signaling domains composed of shuffled signaling motifs. These synthetic cytokine receptors (SCRs) enabled us to study the relationship between signaling domain structure, JAK/STAT signaling, and CAR T cell phenotype. Neural networks and biophysical models trained on the data provide insights that may enable rational design of JAK/STAT signaling domains that promote desired cell function.

## Results

### Signaling motifs induce distinct signaling and T cell phenotype

We created synthetic cytokine receptors (SCRs), chimeric membrane proteins that provide continuous JAK/STAT signaling (Fig. 1A) in the absence of cytokine. SCRs consist of a Put3 coiled-coil domain that induces constitutive dimerization of the erythropoietin receptor (EpoR) transmembrane domain(*36*), IL6 signal transducer JAK-binding box motifs and dileucine internalization motif, and a region containing variable signaling motifs. The fixed regions of the SCR enable continuous signaling such that functional differences between SCR constructs depend only on the variable signaling motifs. SCR transgenes were assembled in a lentiviral cassette (Fig. 1B) encoding the SCR, a viral 2A sequence, and Green Fluorescent Protein (GFP).

**Figure 1.**
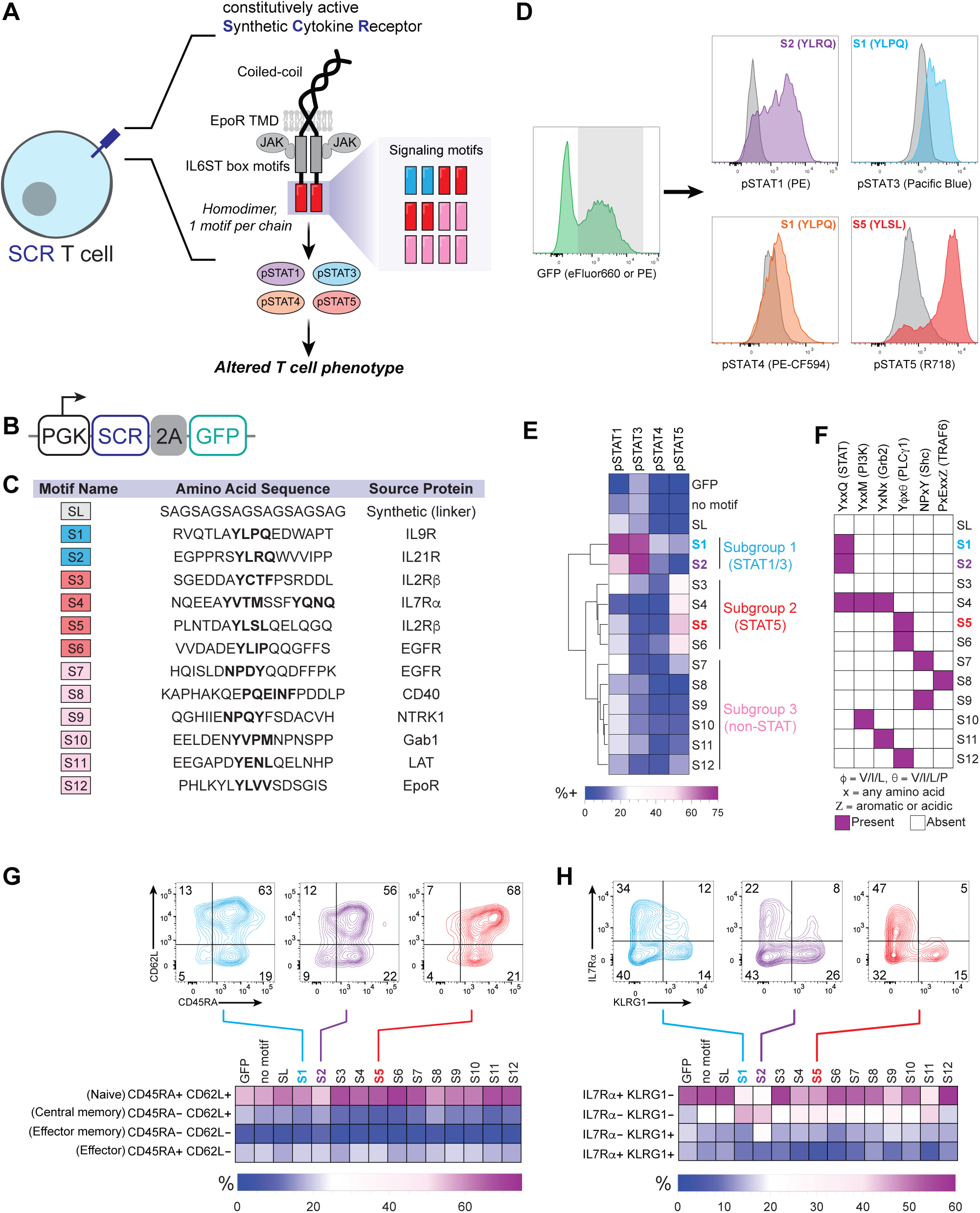
Constitutively active synthetic cytokine receptors (SCR) induce signaling motif-dependent signaling and T cell phenotype. A) Primary human T cells were engineered to express a homodimeric SCR with variable signaling motifs that induce phosphorylation of STATs. SCR, synthetic cytokine receptor; EpoR, erythropoietin receptor; JAK, Janus kinase; IL6ST, interleukin 6 signal transducer; STAT, signal transducer activator of transcription; pSTAT, phosphorylated STAT. B) A lentiviral cassette with a PGK promoter expresses the SCR, a viral 2A sequence, and GFP. PGK, phosphoglycerate kinase; GFP, green fluorescent protein. C) Description of library parts used in the combinatorial library. Each part includes the signaling motif(s) and flanking sequence. Phosphotyrosines are shown in bold. D) Flow cytometry was used to measure phosphorylation of STAT1, STAT3, STAT4, and STAT5 in T cells transduced with SCRs containing varied motifs. E) Quantification of flow cytometry pSTAT measurements for controls and SCRs containing varied motifs. Library parts S1 through S12 were sorted by hierarchical clustering of STAT phosphorylation signature using the Dendrogram function in *Mathematica 13.2*. F) Library parts contain consensus motif sequences with putative binding partners (parentheses) involved in cell signaling. PI3K, phosphatidylinositol-3-kinase; Grb2, growth factor receptor-bound protein 2; PLCy1, phospholipase C gamma 1; Shc, Src homology and collagen protein; TRAF6, tumor necrosis factor receptor–associated factors 6. G) Flow cytometry was used to measure expression of CD45RA and CD62L in T cells transduced with GFP or SCRs containing varied motifs. CD45RA, receptor-type tyrosine-protein phosphatase C; CD62L, L-selectin. H) Flow cytometry was used to measure expression of IL7Ra and KLRG1 in T cells transduced with GFP or SCRs containing varied motifs. IL7Ra, interleukin 7 receptor alpha chain; KLRG1, killer cell lectin-like receptor subfamily G member 1. Values depicted in heatmaps E and G represent mean values of 3 replicates. Values and error bars in J and L represent mean and standard deviation of 3 replicates.

To investigate how individual signaling motifs encode T cell signaling and function, we virally transduced T cells with SCR variants which each contain a distinct signaling motif. The motifs (Fig. 1C) include 12 sequences (S1 through S12) derived from natural signaling proteins, and a synthetic linker sequence (SL). In GFP+ primary human T cells expressing SCRs that each contain one motif, we measured the phosphorylation of STAT1, STAT3, STAT4, and STAT5 (Fig. 1D and E). The percent of cells positive for each phosphorylated STAT varied by motif. The percent of cells positive for each STAT is proportional to the mean fluorescence intensity (fig. S1A). For the remainder of this work, we use percent positivity for pSTAT as a measure of STAT signaling.

Hierarchical clustering of motifs based on STAT phosphorylation revealed three subgroups of motifs (Fig. 1E). Subgroup 1 includes motifs S1 and S2, which induce the strongest phosphorylation of STAT1 and STAT3. Subgroup 2 includes motifs S3 through S6, which induce the strongest phosphorylation of STAT5. Subgroup 3 includes S7 through S12, which induce modest phosphorylation of STATs. Subgroup 3 contains consensus motifs (Fig. 1F) that in their native context bind signaling proteins such as Growth factor receptor-bound protein 2 (Grb2) (S8), Src homology and collagen (Shc) (S9 and S10), Phosphoinositide 3-Kinase (S11), phospholipase C gamma 1 (PLCγ1) (S12)(*8, 37, 38*), or Tumor Necrosis Factor receptor-associated factor 6 (TRAF6) (S7)(*39*). The presence of the consensus motifs and changes in T cell phenotype observed below lead us to suspect SCRs activate signaling pathways associated with these proteins.

Consistent with previous observations by other groups, addition of alanine residues between the transmembrane domain and signaling domain tuned signaling (fig. S1B through D)(*36, 40*). Because SCRs with no added alanines induced the strongest signaling, we used these SCRs for the remainder of experiments. To understand how the number of motif copies in a signaling domain influences signaling, we created SCRs with 1, 2, or 3 copies of motif S1 or S3 (fig. S1E). Adding copies of motif S1 or S3 increased STAT phosphorylation with diminishing returns (fig. S1E through G).

We hypothesized that the motifs, which induce different signaling patterns, would also induce distinct T cell differentiation. We measured expression of differentiation markers receptor-type tyrosine-protein phosphatase C (CD45RA) and L-selectin (CD62L) (Fig. 1G), and expression of longevity marker IL7 Receptor subunit alpha (IL7Rα) and activation marker killer cell lectin receptor G1 (KLRG1) (Fig. 1H). Changes in CD45RA^+^ versus CD62L^+^ were more subtle than changes in IL7Rα^+^ versus KLRG1^-^. The relative size of the undifferentiated IL7Rα^+^ KLRG1^-^ populations induced by S2 and S5 were over 2-fold different, highlighting the different effects of STAT1 and 3 signaing and STAT5 signaling. The overall variation in surface marker expression suggests signaling motifs each contribute differently to T cell differentiation.

### An SCR library with synthetic combinations of signaling motifs generates diverse CAR T cell states

Given the varied T cell signaling and differentiation induced by each motif, we reasoned that SCR signaling domains that contain combinations of these motifs would produce a broad spectrum of function when expressed in CAR T cells. We constructed a library of SCRs with signaling domains that contain random combinations of three to five motifs. To test the effects of these SCRs on CAR T cell function, we co-expressed the SCRs with a second-generation anti-Human Epidermal Growth Factor Receptor 2 (HER2) CAR in primary human T cells (Fig. 2A). Each unique SCR, the anti-HER2 CAR, and GFP were encoded by a single lentiviral vector (Fig. 2B) that was used to transduce the T cells. We transformed the pooled library of SCR-CAR-GFP constructs into *E. coli*, randomly picked 600 colonies for DNA isolation and sequence verification, and created an arrayed library in which each well contains a separate SCR. This arrayed approach allowed us to independently assess the function of each SCR without confounding effects of inter-construct cytokine signaling that would be present in a pooled screen.

**Figure 2.**
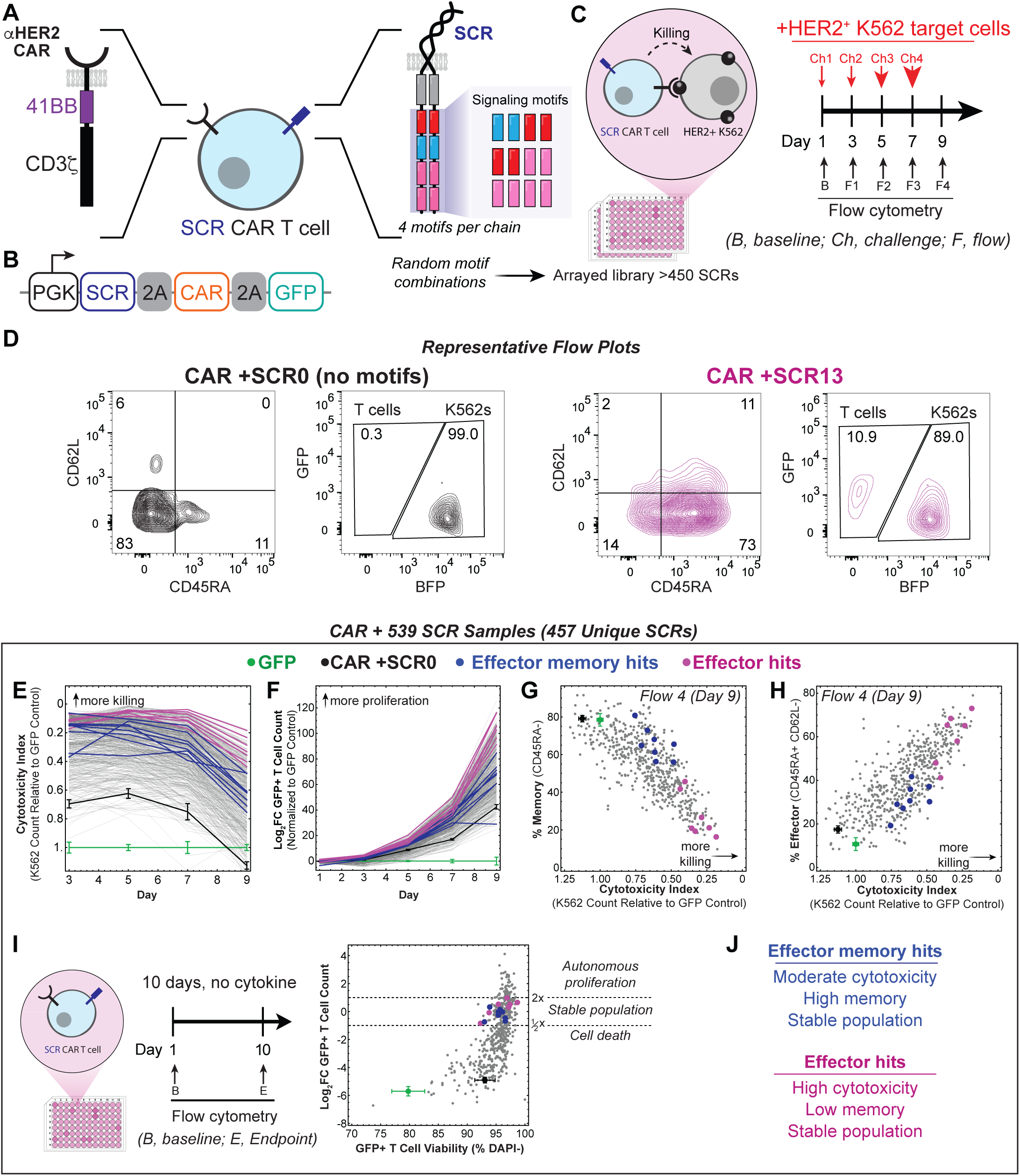
An SCR library with synthetic combinations of signaling motifs generates diverse CAR T cell states. A) Primary human T cells were transduced with the anti-HER2 CAR and SCRs containing an average of four motifs. HER2, Human Epidermal Growth Factor Receptor 2; CAR, chimeric antigen receptor B) A lentiviral cassette with a PGK promoter expresses the SCR, a viral 2A sequence, CAR, viral 2A sequence, and GFP. C) CD4+ and CD8+ SCR CAR T cells with various SCR signaling motif combinations were pulsed four times with increasing numbers of HER2+ K562 target cells. SCR CAR T cell proliferation, differentiation, and cytotoxicity were assessed by flow cytometry at several timepoints. D) Representative flow cytometry plots of SCR CAR T cells and HER2+ K562 target cells on day 9 of co-culture. E-H) SCR CAR T cell cytotoxicity, proliferation, memory (CD45RA-), effector (CD45RA+ CD62L-), and cytotoxicity were assessed by flow cytometry over the 9-day co-culture with HER2+ K562 cells. Cytotoxicity (E) and GFP+ T cell count (F) of SCR CAR T cells and controls were measured on days 3, 5, 7, and 9. Cytotoxicity of SCR CAR T cells and controls versus memory (G) or effector (H) T cell population. I) SCR CAR T cells with various SCR signaling motif combinations were cultured in the absence of exogenous cytokine for 10 days. GFP+ cell count was measured on days 1 and 10 by flow cytometry. GFP+ cell survival (DAPI-) was measured on day 10 by flow cytometry. CAR samples in E and F were transduced with an anti-HER2 CAR, an SCR with no signaling motifs, and GFP. GFP samples in E and F were transduced with GFP. DAPI, 4’,6-diamidino-2-phenylindole. J) Description of effector memory hits and effector hits identified from arrayed screen. Example effector memory hits (blue) and effector hits (magenta) are shown in E through I. Values and error bars for CAR and GFP samples in E through F represent mean and standard deviation of 14 samples spread across seven 96-well plates.

We repeatedly pulsed SCR CAR T cells with increasing doses of engineered HER2+ K562 target cells and performed flow cytometry measurements at various timepoints to assess the co-culture (Fig. 2C). T cell differentiation was assessed by expression of CD45RA and CD62L (Fig. 2D and fig. S2A through D). Proliferation was assessed by fold change in GFP+ T cell count and cytotoxicity was assessed by reduction in K562 cell count. Nearly all SCRs increased CAR T cell cytotoxicity (Fig. 2E) and proliferation (Fig. 2F) relative to control T cells expressing only CAR. These SCR-dependent increases were maintained for the duration of the co-culture experiment. Following the final pulse of target cells, which contains the largest number of target cells, the cytotoxicity of the CAR control (expressing an SCR with no motifs, SCR0) was indistinguishable from that of control T cells expressing only GFP. At the end of the co-culture, the T cell cytotoxicity was negatively correlated with T cell memory population (Fig. 2G) and positively correlated with T cell effector population (Fig. 2H). These correlations were present on day 5 (fig. S2E) and more pronounced on day 9 (fig. S2F). Together, these data suggest the existence of a tradeoff between CAR T cell memory formation and cytotoxicity. This tradeoff can be tuned by engineering CAR T cell JAK/STAT signaling.

To test for the ability of SCRs to provide T cell survival signals, we cultured SCR CAR T cells in the absence of cytokine for 10 days (fig. 2I). We performed flow cytometry on day 1 and day 10 to measure GFP+ T cell count and the percent of GFP+ T cells that were live (DAPI-). SCRs had varied effects on CAR T cell survival. Some SCRs promoted T cell survival and resulted in stable populations (final population cell count between 0.5x and 2x initial cell count) over the 10-day culture. Other SCRs promoted autonomous CAR T cell proliferation or provided insufficient survival signals to prevent T cell death.

We identified two groups of SCRs that increase CAR T cell cytotoxicity and proliferation and enabled survival of a stable CAR T cell population in the absence of cytokine (fig. 2J). The first group contains effector memory hits characterized by high memory and moderate cytotoxicity (Fig. 2G) with a stable population (Fig. 2I). At the end of the co-culture, most of these cells were in the effector memory (CD45RA-CD62L-) state (fig. S2D). The second group contains effector hits characterized by low memory and high cytotoxicity (Fig. 2G) with a stable population (Fig. 2I). At the end of the co-culture, most of these cells were in the effector (CD45RA+ CD62L-) or naïve-like (CD45RA+ CD62L+) state (Fig. 2D). Upon stimulation, the effector hits proliferated more than effector memory hits (Fig. 2F). The results of this screen demonstrate that SCRs with synthetic signaling domains composed of recombined signaling motifs can steer CAR T cells to diverse phenotypic states with a spectrum of *in vitro* anti-tumor activity.

### SCRs with synthetic signaling domains improve CAR T cell anti-tumor activity

We further characterized the signaling and function of an effector hit (SCR421) and a memory effector hit (SCR516) (Fig. 3A). Data for several other SCR hits of interest (fig. S3A) can be found in figure S3. The chosen SCR hits had distinct activity in the arrayed screen (fig. S3B). The *in vitro* anti-tumor activity of SCR421 and SC516 were benchmarked against two SCRs with signaling domains derived from the IL7 receptor alpha chain (IL7Rα) and the IL21 receptor, which are known to improve CAR T cell efficacy(*13, 19, 41*). We constructed SCRs containing residues P329 to Q549 of IL7Rα or R275 to S538 of IL21R, removing the IL7Rα and IL21R box motifs but preserving downstream signaling motifs.

**Figure 3.**
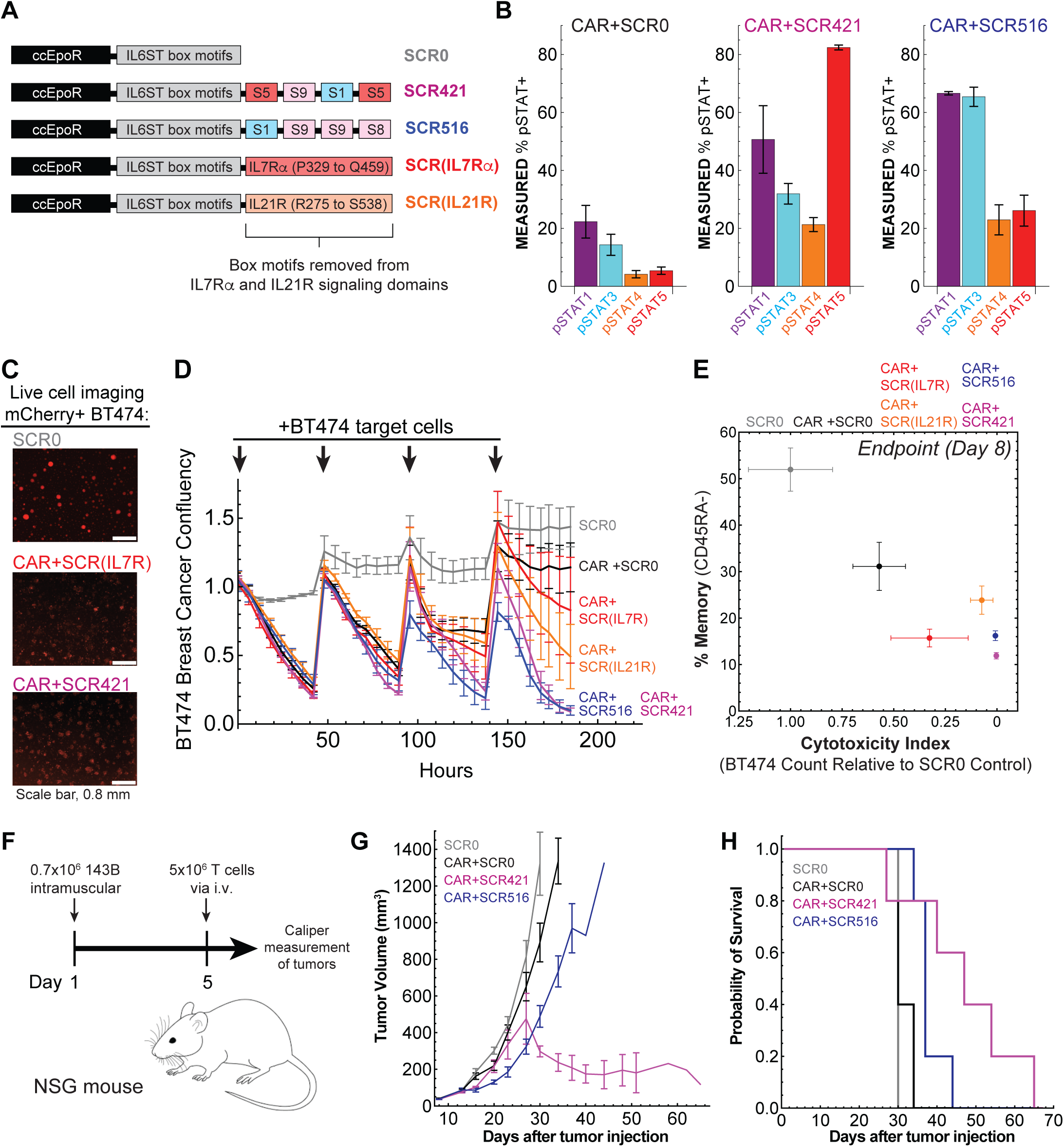
SCRs with synthetic signaling domains improve CAR T cell anti-tumor activity. A) Schematics of SCR0, SCR421, SCR516, SCR(IL7Ra) and SCR(IL21R). B) Quantification of the percent of SCR CAR T cells positive for pSTAT1, 3, 4, or 5. C) Live cell images of SCR CAR T cells co-cultured with mCherry+ BT474 breast cancer cells at the end of an 8-day co-culture. D) SCR CAR T cells were pulsed with mCherry+ BT474 breast cancer cells at 0, 48, 96, and 144 hours. Confluency of BT474 cells was monitored by live cell imaging to assess SCR CAR T cell cytotoxicity. Confluency was normalized to average confluency of SCR0 samples at time 0. E) Flow cytometry was used to assess SCR CAR T cell cytotoxicity and memory on day 8 of co-culture with BT474 cells. F-H) NSG mice were injected with 0.7×10^6^ 143B on day 1 and 5×10^6^ T cells on day 5 (F). Tumor volumes were measured two to three times weekly by caliper (G). Survival of mice was monitored for the duration of the experiment (H). n = 5 mice per group. Values and error bars in B, D, E, and G represent mean and standard error of 3-5 replicates.

We used flow cytometry to measure the phosphorylation of STAT1, 3, 4, and 5 in resting CAR T cells expressing SCR0, SCR421, and SCR516 (Fig. 3B). The percent of cells positive for each pSTAT varied by SCR. SCR0, which contains no signaling motifs aside from the box motifs, displayed the lowest STAT phosphorylation. SCR421 induced high pSTAT5, moderate pSTAT1, and low pSTAT3 and pSTAT4. The strong STAT5 signal is consistent with the presence of two copies of the S5 motif, which induces the most pSTAT5 of the 13 motifs in the library. SCR516 induced high pSTAT1 and pSTAT3, and low pSTAT4 and pSTAT5. These results are consistent with prior observations that pSTAT5 is required for T cell effector function and pSTAT3 plays a critical role in maintaining T cell memory.

To investigate the *in vitro* cytotoxicity and differentiation of SCR421 and SCR516, we co-cultured SCR CAR T cells with mCherry+ BT474 breast cancer cells added in serial pulses. We used live cell imaging (Fig. 3C and D, fig. S3C) to monitor killing of BT474 cells throughout the co-culture and flow cytometry (Fig. 3E) to monitor killing of BT474 cells and CAR T cell differentiation at the end of the co-culture. Following the first BT474 pulse, all SCR CAR T cell samples induced BT474 death (Fig. 3D). The killing of BT474 cells diverged over subsequent pulses. At the end of the co-culture, CAR T cells with SCR421 and SCR516 displayed more killing than CAR T cells with SCRs containing the IL7Rα or IL21R signaling domains.

These live cell imaging observations are consistent with killing measurements made by flow cytometry (Fig. 3E). As previously observed in the co-culture of SCR CAR T cells and K562 cells, memory and cytotoxicity were anti-correlated. However, in contrast to the co-culture with K562 cells, SCR421 and SCR516 induced similar memory during co-culture with BT474 cells. Despite inducing lower memory than SCRs with IL7Rα or IL21R signaling domains, SCR421 and SCR516 maintained CAR T cell cytotoxicity throughout the co-culture. Taken together, these results demonstrate that recombination of signaling motifs can produce novel signaling domains with *in vitro* anti-tumor activity superior to that of natural signaling domains.

Next, we assessed the anti-tumor activity of SCR421 and SCR516 against the 143B osteosarcoma model in NOD.Cg-*Prkdc^scid^ Il2rg^tm1Wjl^*/SzJ (NSG) mice (Fig. 3F through H, fig. S3D). T cells expressing SCR0 or CAR and SCR0 offered minimal control of 143B tumor growth (Fig. 3G). CAR T cells expressing SCR516 significantly slowed growth of 143B tumors relative to CAR T cells expressing SCR0. SCR421 induced the strongest anti-tumor activity, resulting in reduction of tumor volume. Both SCR421 and SCR516 extended the survival of mice relative to SCR0 (Fig. 3H). Notably, SCR27, which contains four S5 STAT5-binding motifs, behaved similarly to SCR421 *in vitro* and resulted in reduction of tumor volume *in vivo* (fig. S3D), but failed to extend mouse survival. Previous work demonstrated constitutive activation of STAT5 can protect against Tox-induced exhaustion and loss of T cell anti-tumor efficacy(*15–17*). These prior observations may explain why CAR T cells expressing SCR421 and SCR27, which have strong STAT5 signaling, have potent anti-tumor efficacy despite differentiating to the effector state. Relative to SCR27, SCR421 has similar STAT5 signaling and increased STAT1, STAT3, and STAT4 signaling. SCR421 also contains an Shc-binding motif that later analyses (Fig. 4D and E) suggest contributes to anti-tumor function. The superior *in vivo* anti-tumor activity of SCR421 relative to SCR27 underscores the utility of motif recombination that enables customization of signaling programs. Our results demonstrate that SCRs with synthetic signaling domains can drive CAR T cells to distinct states with increased *in vivo* anti-tumor efficacy.

**Figure 4.**
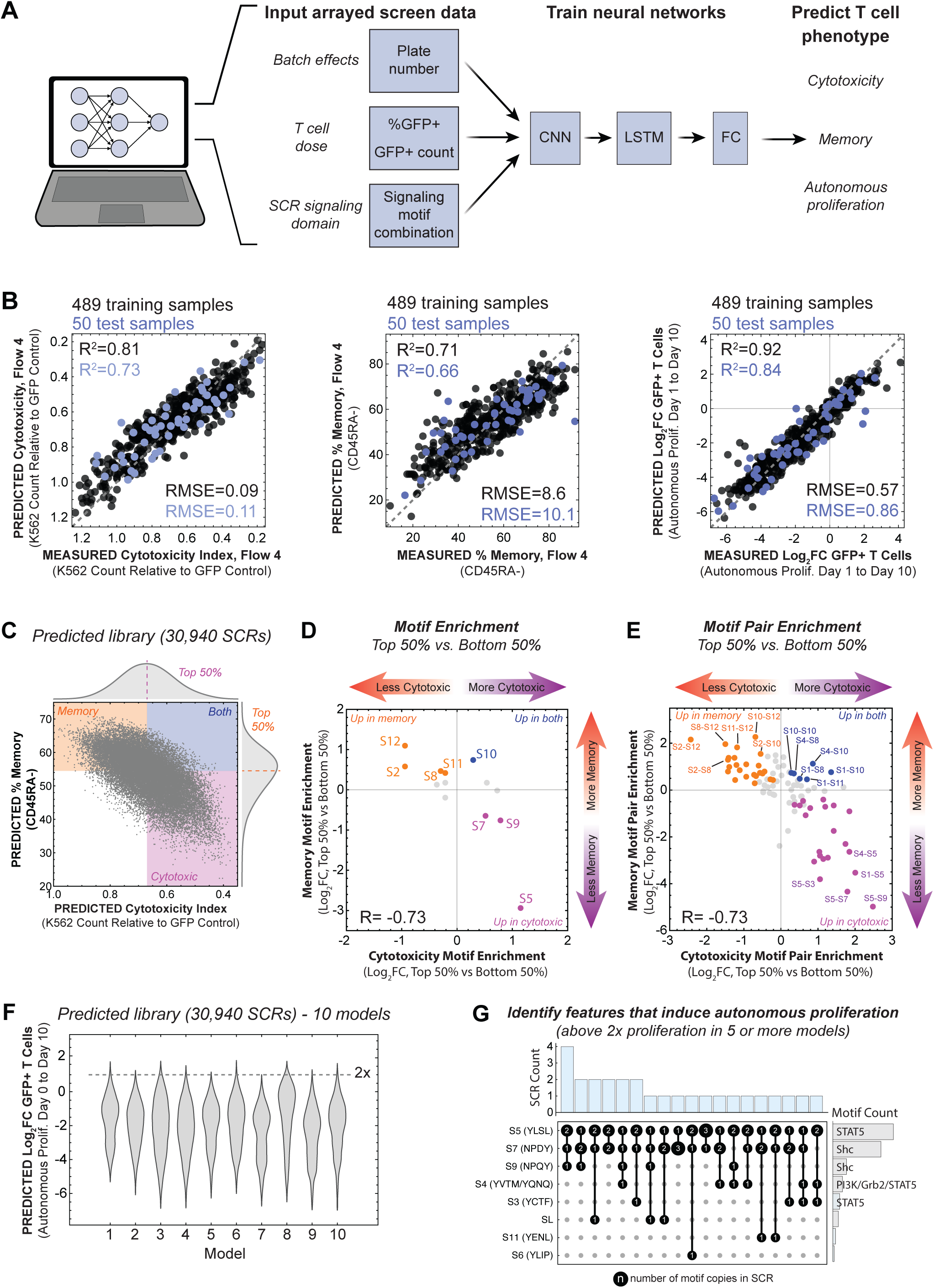
Neural networks predict CAR T cell memory, cytotoxicity, and autonomous proliferation encoded by SCR signaling motif combinations. A) Schematic of the neural network used to predict SCR CAR T cell phenotype. CNN, convolution neural network; LSTM, long short-term memory; FC, fully connected. B) Neural networks trained on array data predict the cytotoxicity, memory, and unstimulated proliferation of SCR CAR T cells in the training sets (black) and the withheld test sets (magenta). Gray dashed lines represent y = x. RMSE, root mean squared error. C) Trained neural networks were used to predict the cytotoxicity and memory encoded by 30940 SCRs containing one to four variable signaling motifs. Marginal distributions for cytotoxicity and memory are projected on the x and y axes. Dashed lines demarcate the top 50% and bottom 50% of samples for cytotoxicity and memory. Predictions represent the mean for n = 10 neural networks with different hyperparameters. D,E) Neural network predictions of motif and motif pair enrichment cytotoxicity and memory. Plots depict Log2FC for motifs and motif pair frequencies of SCRs in top 50% and bottom 50% for each phenotype. Motifs and motif pairs with statistically significant enrichment (*p*<0.05) are shown in orange, magenta, or blue. Each *p*-value was calculated by performing a Student’s t-test to compare the observed FC values from 10 model predictions to the 10 expected FC values for the null hypothesis. FC, fold change. F) Distributions of SCR CAR T cell unstimulated proliferation for 30940 SCRs predicted by 10 neural networks trained on arrayed screen data. Gray dashed line represents threshold for SCR CAR T cell samples predicted to proliferate two-fold in 10 days. G) UpSet plot showing composition of SCRs predicted to induce autonomous proliferation in at least five neural network models. For each SCR motif combination (blue bars), the counts of motifs in the SCR are shown in black circles. Total motif counts across all SCRs are depicted to the right (gray bars).

### Neural networks predict CAR T cell memory, cytotoxicity, and autonomous proliferation encoded by SCR signaling motif combinations

Experimental results (Figs. 1 and 2) demonstrate that motif recombination in JAK/STAT signaling proteins can generate a diversity of CAR T cell function. To understand how signaling motifs in the SCR signaling domain encode CAR T cell function, we used the screening data (Fig. 2) to train neural networks that predict CAR T cell cytotoxicity, memory, and autonomous proliferation (Fig. 4A). The neural networks use the independent variables of the screening data as inputs. These independent variables include the 96-well plate in which each sample was located, the GFP+ T cell count and percent of T cells that were GFP+ in the baseline flow cytometry measurement, and identity of the signaling motifs in the SCR signaling domain. These independent variables account for plate-dependent batch effects, CAR T cell dose, and signaling domain structure. The independent variables are pre-processed (using embedding or fully connected layers with linear activation), concatenated, and passed through a convolutional layer, a long short-term memory layer, and fully connected layers to produce quantitative predictions of cytotoxicity, memory, and autonomous proliferation. A detailed description of the neural network architecture and training can be found in the **Materials and Methods** section.

Neural networks were trained on 489 training points using k-fold averaged cross validation to select hyperparameters. For each T cell phenotype, we obtained an ensemble model consisting of the ten best models found during k-fold cross validation. A test set of 50 points was withheld during training and used to assess model accuracy. Experimentally measured and neural network predicted values for training and test sets demonstrate that neural networks can predict CAR T cell cytotoxicity, memory, and autonomous proliferation (Fig. 4B) encoded by SCR signaling motifs.

We used the trained neural networks to predict the cytotoxicity, memory, and autonomous proliferation of 30,940 SCRs containing one, two, three, or four of the 13 signaling motifs in our library in all possible combinations and arrangements. To remove batch effects and effects of CAR T cell dose, predictions were made using values that correspond to measurements made in a single plate (plate 1) and uniform GFP+ T cell count (5869 cells) and percent GFP+ T cells (67%). The cytotoxicity and memory of SCR CAR T cells in the predicted library (Fig. 4C) recapitulate the tradeoff between cytotoxicity and memory observed in experimental data (Fig. 2G). Marginal distributions of cytotoxicity and memory for the simulated library are shown on the x and y axes. SCR CAR T cell samples predicted to be in the top 50% of cytotoxicity (magenta), memory (orange), or both (blue) are shown in colored boxes. The number of SCR CAR T cell samples predicted to be in the top 50% of both memory and cytotoxicity (4193, 13.5%) is less than the number expected (7735, 25%) in the case of uncorrelated memory and cytotoxicity.

To identify motifs or motif pairs that encode memory, cytotoxicity, or both, we performed enrichment analysis on the predicted library data (Fig. 4D and E). We calculated the predicted Log2 fold change (FC) in motif abundance between the top 50% and bottom 50% for cytotoxicity and memory. Motifs with enrichment values greater than zero promote the T cell phenotype of interest relative to motifs with enrichment values less than zero. Motifs that promote cytotoxicity (magenta), memory (orange), or both (blue) with statistically significant (*p* < 0.05) enrichment or de-enrichment in both cytotoxicity and memory are shown in color. We performed a similar enrichment analysis for motif pairs (Fig. 4E). For both motifs and motif pairs, enrichment in cytotoxicity and memory were anti-correlated (R = −0.73). This indicates that at least some of the tradeoff between cytotoxicity and memory is encoded in the signaling motif combination. Motif S5, which has a YLSL motif that strongly activates STAT5, was the most enriched in the cytotoxic samples. Motifs S7 and S9, which each have an NPxY motif that binds Shc, are also enriched in cytotoxic samples. Motifs S12, S2, S8, and S11 are enriched in the memory samples. These four motifs have distinct STAT signaling patterns (Fig. 1E) and diverse consensus motifs (Fig. 1F). Only motif S10, the PI3K-binding YVPM motif from Grb2-associated binder 1 (Gab1), was significantly enriched in both cytotoxic and memory samples.

Many motif pairs were significantly enriched in the memory samples, cytotoxic samples, or both. The enrichment magnitudes of motif pairs were larger than the enrichment magnitudes of motifs. All five of the motif pairs most enriched in cytotoxic samples contain motif S5, suggesting that S5 and pSTAT5 are potent inducers of cytotoxicity. Moreover, two of the motif pairs most enriched in cytotoxicity, S1-S5 and S5-S9, are found in SCR421. This suggests these two motif pairs are responsible for much of the cytotoxicity induced by SCR421. One of the motif pairs most enriched in both cytotoxicity and memory, S1-S8, is found in SCR516. The S1-S8 motif pair may encode much of the balanced phenotype induced by SCR516. These analyses underscore the utility of machine learning modeling to identify motifs that drive desired cell functions.

Previous research on constitutively active chimeric cytokine receptors indicated some signaling domains can induce dangerous autonomous proliferation in CAR T cells. For example, Zip7R, the ZipR with an IL7R signaling domain, induced deadly proliferation in mice(*30*). In the arrayed screen we observed autonomous proliferation of CAR T cells driven by a subset of SCRs (Fig. 2I). We sought to use models trained on the autonomous proliferation data to identify signaling motifs that induce autonomous proliferation. We used the ensemble of 10 models trained during k-fold cross validation to identify SCRs predicted to drive 2-fold or greater autonomous CAR T cell proliferation over 10 days in at least 5 of the models (Fig. 4F). An UpSet plot of these SCRs (Fig. 4G) shows that motifs S5, S7, S9, S4, and S3 cooperate to induce autonomous proliferation. All the identified SCRs contain S5, which activates pSTAT5. Nearly all these SCRs contain S9 or S10, which contain NPxY consensus motifs that binds Shc, leading to subsequent activation of Grb2. The only SCR without an S7 or S9 motif contains S4, which has binding sites for Grb2 and PI3K. Motif S4 is derived from the IL7Rα chain, indicating it may be involved in autonomous proliferation of CAR T cells expressing Zip7R. Our machine learning-based analysis suggests a core group of mitogenic signals encompassing pSTAT5, Grb2, and PI3K drive autonomous proliferation. Prior research has demonstrated that these signals can combine to induce malignant cell behavior. These findings demonstrate models can be used to predict signaling domain structural features that drive undesired cell functions like autonomous proliferation. Such models may enable design of safer synthetic receptors and CAR T cells.

### A continuous landscape of JAK/STAT signaling encodes a corresponding landscape of CAR T cell phenotype

Our results thus far show that signaling motifs each induce distinct signaling and that signaling motif recombination generates a variety of CAR T cell phenotypes. To further investigate, we combined experimental data and modeling to quantitatively link signaling domain structure, cell signaling, and CAR T cell phenotype. In the arrayed screen of SCR CAR T cells and K562 cells, we observed a negative correlation between T cell cytotoxicity and memory (Fig. 2G). In the simulated library, we observe this negative correlation and a Pareto front (Fig. 5A and B). We hypothesized that signaling changes gradually from the first SCR along the Pareto front, which induces maximum memory, to the last SCR along the Pareto front, which induces maximum cytotoxicity. To understand how cell signaling encodes the trade-off between memory and cytotoxicity (Fig. 5C), we set out to map the signaling in the 476 SCRs along the Pareto front.

**Figure 5.**
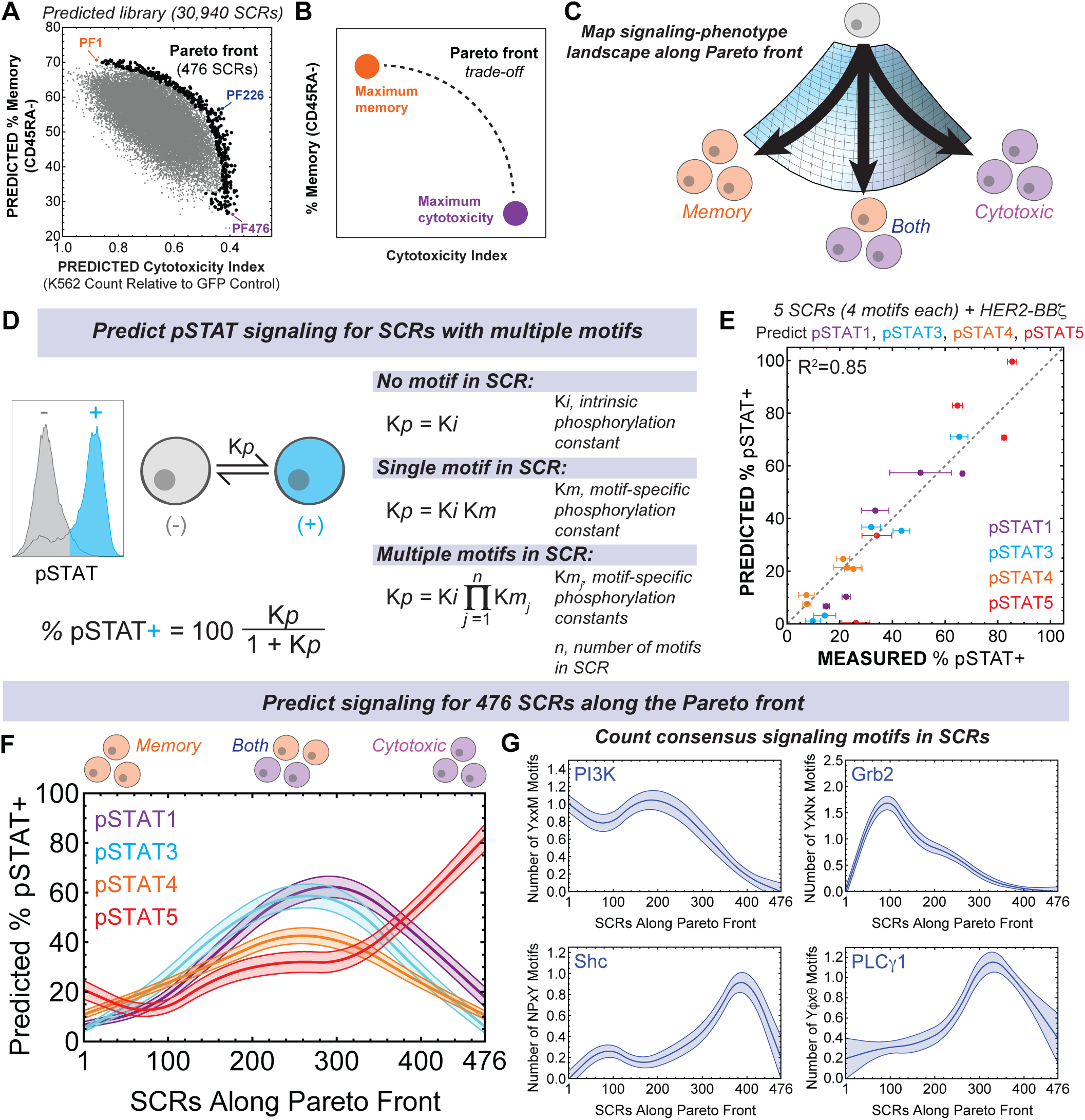
A continuous landscape of JAK/STAT signaling encodes a corresponding landscape of CAR T cell phenotype. A) Trained neural networks were used to predict the cytotoxicity and memory encoded by 30940 SCRs containing one to four variable signaling motifs. The SCR library data suggests a trade-off between memory and cytotoxicity. 476 SCRs near the Pareto front are marked in black. B) Schematic of a Pareto front. C) We combined flow cytometry measurements of SCR pSTAT signaling, flow cytometry measurements of SCR CAR T cell cytotoxicity and memory, neural network predictions, and biophysical models to map the signaling-phenotype landscape of SCR CAR T cells. D) Description of a biophysical model to predict the percent of T cells positive for pSTAT. E) Measured and predicted percent of SCR CAR T cells positive for pSTAT1, 3, 4, and 5. To make predictions the model in D was fit to measurements of SCR-induced pSTAT signaling (Fig. 1E and fig. S5A) to determine K*i* and K*m* values. Fitted parameters include four K*i* values: one each for pSTATs 1, 3, 4, and 5. We determined four K*m* values for each signaling motif (SL and S1 through S12). K*m* values were calculated as –y/(K*i*(−100+y)) where y is the percent of cells positive for pSTAT. Measured and predicted values for pSTAT signaling for SCRs with no signaling motifs (fig. S5B) and SCRs with 0, 1, 2, or 3 copies of signaling motifs (fig. S5C) can be found in the supplement. Model fitting and subsequent predictions were performed in *Wolfram Mathematica 14.1*. F) The fitted model of SCR-induced pSTAT signaling was used to predict the pSTAT1, 3, 4, and 5 signaling for the 476 SCRs along the Pareto front in Fig. 5A. G) Count of consensus signaling motifs suspected to bind PI3K, Grb2, Shc, and PLCγ1 for the 476 SCRs along the Pareto front in Fig. 5A. Values and bands in F and G represent the mean and standard error for predicted signaling.

To map SCR signaling to CAR T cell cytotoxicity and memory, we used the flow cytometry phosphorylation measurements of T cells expressing SCRs with single motifs (Fig. 1E) or multiple motifs (fig. S5A) to fit a model (Fig. 5D) that predicts STAT signaling induced by different SCR motif combinations. For each pSTAT, cells exist in one of two states: negative for pSTAT and positive for pSTAT. The equilibrium between the negative and positive states is defined by the equilibrium K*_p_*. The percent of cells in the positive state is 100*K*_p_*/(1+K*_p_*). In the absence of cytokine or SCR signaling K*_p_* is equal to K*_i_*, the intrinsic equilibrium constant for phosphorylation. Upon addition of signaling motifs to the SCR, K*_p_* can be described by K*_i_*\*K*_m_*, where K*_m_* is a motif-specific phosphorylation constant. For each motif in our library, the K*_m_* for pSTAT1, 3, 4, and 5 can be calculated from the phosphorylation data in Figure 1E. For SCRs with more than one motif, K*_p_* is equal to K*_i_* times the product of the K*_m_* for each motif in the SCR.

The agreement between measured STAT signaling and predicted STAT signaling for five SCRs (R^2^ = 0.85) is shown in Figure 5E. The fitted K*_i_* values for pSTAT1, 3, 4, and 5 produce predictions of intrinsic STAT signaling similar to measured STAT signaling induced by SCR0 (fig. S5B). This model also predicts the STAT3 or STAT5 signaling (fig. S5C) for SCRs with multiple copies of motif S1 or S3 (fig. S1G). Data for SCR0 or SCRs with multiple copies of motifs S1 or S3 were not used to fit the model, thus agreement between the measured and predicted values indicates the model has predictive power. Based on the agreement between the data and model, we used the model to predict STAT signaling for SCRs along the Pareto front (Fig. 5F). The predicted percent of CAR T cells positive for each pSTAT changes gradually from SCRs PF1 through PF476 along Pareto front. The overall trend of the predicted signaling is similar to the signaling calculated by summing the signaling of the individual motifs in the SCR (fig. S5D). The predictions of the fitted model have realistic magnitudes not exceeding 100%.

SCRs that induce high memory and low cytotoxicity are predicted to have little STAT signaling, similar to SCR0. In SCRs with balanced memory and cytotoxicity (example: Fig. 5A, PF226), the predicted signaling is dominated by STAT1 and 3, with lower STAT4 and 5 signaling. Similar signaling was measured for SCR516 and SCR548, which have balanced memory and cytotoxicity (fig. S5A). In the transition to cells with low memory and high cytotoxicity, the predicted STAT1, 3, and 4 signaling decrease while STAT5 signaling increases. In the most cytotoxic samples, predicted signaling is dominated by STAT5 with low STAT1, 3, and 4 signaling. This predicted signaling is similar to that of the effector hits SCR421 and SCR27 (fig. S5A). We also mapped signaling along the diagonal (fig. S5I) and the edge away from the Pareto front (fig. S5J) of our predicted library. These signaling trajectories also show gradual changes in STAT signaling as CAR T cell phenotype changes. The model is consistent with observations that pSTAT3 is required for T cell memory and pSTAT5 is required for T cell effector function.

Maps of STAT1, STAT3, STAT4, and STAT5 signaling in SCRs throughout the predicted library (fig. S5K) demonstrate the tuning of STAT signaling can be used to access a continuous spectrum of CAR T cell cytotoxicity and memory. While STAT signaling is critical for determining T cell function, motifs in our library likely activate other signaling pathways. The average count of consensus signaling motifs that bind PI3K, Grb2, Shc, and PLCγ1 also varies the Pareto front (Fig. 5G). Motifs that bind PI3K (YxxM) and Grb2 (YxNx) are more common in SCRs that induce CAR T memory. Motifs that bind Shc (NPxY) or PLCγ1 (Yϕxθ) are more common in SCRs that induce cytotoxicity. These results indicate that changes in cell signaling create a continuous spectrum of CAR T cell cytotoxicity and memory and encode the trade-off between these two phenotypes.

## Conclusions

We created a library of synthetic JAK/STAT signaling proteins with novel motif combinations and tested hundreds of these proteins for their effects on CAR T cell survival, proliferation, memory formation, and cytotoxicity. The *in vitro* and *in vivo* data, along with models trained on the data, yield insights into the connection between signaling domain structure, cell signaling, and CAR T cell phenotype. Synthetic signaling domains generated by recombination of a small number of signaling motifs can induce a wde range of cytotoxicity, differentiation, survival, and proliferation. Nearly all synthetic signaling domains increased CAR T cell proliferation and *in vitro* killing of target cells.

We identified several SCRs with synthetic signaling domains that improved *in vitro* and *in vivo* anti-tumor efficacy without dangerous autonomous proliferation. These findings agree with prior observations about the role of JAK/STAT signaling in T cell biology. For example, we and others observe that signaling dominated by STAT3 and STAT1 yielded balanced CAR T cell memory and cytotoxicity that improves anti-tumor activity(*42, 43*). Signaling dominated by STAT5 drives potent CAR T cell effector function and protects against loss of anti-tumor activity(*17, 44*). We also observed evidence that pSTAT5 cooperates with mitogenic signals such as Grb2 (downstream of Shc) and PI3K to induce autonomous cell proliferation(*45*).

We explored the relationship between structure, signaling, and function without considering complicating variables present in natural cytokine signaling. Our use of constitutively dimerized SCRs with invariable box motifs and a limited set of signaling motifs removes potential effects that might arise in natural cytokine signaling systems. These kinetic effects might arise from differences in cytokine concentration(*46*), receptor trafficking kinetics(*47*), or differences in STAT phosphorylation due to unique JAKs recruited by various box motifs(*48*). Extensions of this work may need to consider these parameters.

The screening and machine learning approach used here to study CAR T cell biology can be extended to include other immune cell biology (cell types or phenotypes) of interest. For example, one might perform screens to measure CAR T cell engraftment(*49*), macrophage polarization and phagocytosis, or natural killer cell survival encoded by a library of synthetic JAK/STAT signaling domains. Porting synthetic signaling domains into chimeric cytokine receptors that require a cytokine to dimerize would greatly expand the JAK/STAT signaling space and provide a new tool kit to steer cells to desired fates. We performed our measurements of SCR-driven cell survival and autonomous proliferation in the absence of exogenous cytokine and other signals that CAR T cells might encounter. More sophisticated *in vitro* screens or *in vivo* screens may accurately identify SCRs that have low risk of autonomous proliferation. Such data could then be used to train models that provide insights into engineering more effective immune cell therapies.

The choice of signaling motif determines the binding partners for receptor signaling domains, thereby influencing cell signaling and cell phenotype. Our results suggest that cells may act as state machines that toggle between unphosphorylated and phosphorylated states. The equilibrium between these two states is set by the motifs in actively signaling receptors, wherein each motif contributes free energy to the phosphorylation of STATs and likely other signaling proteins. Variations in the phosphorylation equilibria for different STATs create a spectrum of STAT signaling. Our work provides evidence that STAT-dependent cell phenotype is encoded by the absolute and relative amounts of STAT signaling. Rather than specific combinations of STAT signals encoding discrete phenotypic states, a continuum of signaling defines a corresponding continuum of phenotypes. The tradeoff between CAR T cell memory and cytotoxicity is also encoded by this continuum of signaling such that the relative memory and cytotoxicity change continuously as cell signaling changes. Thus, CAR T cells are tunable state machines in which anti-tumor phenotypes can be fine-tuned by recombining motifs in receptor signaling domains.

## MATERIALS AND METHODS

### Gene synthesis and viral vector construction

DNA fragments encoding the variable library parts (SL, S1 through S12), SCR, P2A-GFP, and P2A-CAR were codon optimized for expression in human cells using ThermoFisher GeneArt’s website tool and synthesized by ThermoFisher GeneArt. The SCR is a chimeric protein consisting of a signal peptide, myc tag, Put3 coiled-coil domain, erythropoietin receptor transmembrane domain, gp130 box motifs and disordered region (residue 642 to residue 725), and gp130 dileucine internalization motif (residue 780 to residue 796). Additional versions of the SCR with reduced magnitude of signaling were created by adding 1, 2, or 3 alanine residues between the transmembrane domain and gp130 fragment. One non-functional SCR was created by mutating the box motif residues IWPNVPDPS to IGSNVPDGS and the residues DVSVVEIEAN to DGSGSEIEAN, replacing critical residues for JAK binding. The P2A-CAR is a chimeric protein consisting of a P2A sequence, signal peptide, hemagglutinin (HA) tag, anti-HER2 4D5 single chain variable fragment, CD8a hinge and transmembrane domain, 41BB-derived costimulatory domain, and CD3z signaling domain.

To create a lentiviral vector containing SCR-P2A-GFP, a pHR lentiviral vector containing a PGK promoter (Addgene #79120) was BamHI-digested and gel (Invitrogen UltraPure Agarose #16500500) purified using a NucleoSpin Gel and PCR Clean-up Kit (Macherey-Nagel #740609.50). Linearized plasmid, a fragment encoding SCR, and a fragment encoding P2A-GFP were assembled by In-Fusion cloning (Takara Bio USA #638949).

For variable library parts and P2A-CAR, synthesized DNA was subcloned by ThermoFisher into standard pMA or pMX plasmid vectors. To obtain DNA fragments for assembly, pMA and pMX vectors were BfuAI-digested and fragments were isolated by agarose gel purification using a NucleoSpin Gel and PCR Clean-up Kit. For combinatorial library assembly, variable part fragments were pooled and used in subsequent In-Fusion reactions into BamHI-digested pHR-PGK-SCR-P2A-GFP. For targeted construct assembly, variable part fragments were added in separate In-Fusion reactions to pHR-PGK-SCR-P2A-GFP. To add CAR, pHR-PGK-SCR-P2A-GFP plasmid already containing desired motifs was BamHI-digested and P2A-CAR was added by In-Fusion cloning. All constructs were sequence verified before use. Amino acid sequences for variable parts are listed in Figure 1 (e.g., S1: RVQTLAYLPQEDWAPT).

### Primary human T cell isolation and culture

Primary CD4+ and CD8+ T cells were isolated from blood of anonymous donors by negative selection using a human T cell isolation kit (STEMCELL Technologies #17951). T cells were cryopreserved in RPMI 1640 plus GlutaMAX (Gibco #61870036) with 20% human AB serum (Sigma-Aldrich, #H5667-100ml) and 10% DMSO (Santa Cruz Biotechnology #SC-358801). Upon thawing, T cells were cultured in Immunocult-XF T Cell Expansion medium (STEMCELL Technologies #10981) supplemented with 30 units/mL IL-2 (STEMCELL Technologies #78145). Before co-culture with cancer cells, T cells were transferred to Immunocult-XF T Cell Expansion medium without IL-2.

### Lentiviral transduction and sorting of human T cells

Pantropic vesicular stomatitis virus G (VSV-G) pseudotyped lentivirus was produced via transfection of LentiX 293T cells (Takara #632180) with a pHR’SIN:CSW transgene expression vector and the viral packaging plasmids pCMVdR8.91 and pMD2.G using FuGENE HD (Promega, #E2312). Primary T cells were thawed the same day and after 24 hours in culture, were activated with ImmunoCult Human CD3/CD28 T Cell Activator cocktail (STEMCELL Technologies #10991) at 25 μL per 1 × 10^6^ T cells. At 48 hours and 72 hours, viral supernatant was harvested and activated primary human T cells were exposed to the virus for 48 hours in 6-well or 24-well plates (mouse experiments, phosphoproteomics) or in 96-well plates (arrayed screens). For arrayed experiments, no additional T Cell Activator cocktail was added during or after T cell exposure to virus. For mouse and phosphoproteomics samples, additional T Cell Activator cocktail was added at 25 μL per 1 × 10^6^ T cells upon addition of virus and upon replacing viral supernatant with fresh Immunocult-XF T Cell Expansion medium. Activated T cells were allowed 6-10 days without T Cell Activator cocktail to rest before subsequent experiments. For arrayed experiments and mouse experiments, T cells were not sorted. For phosphoproteomics experiments, GFP+ T cells were sorted using a FACS Aria II approximately 6 days after initial T cell activation.

### Cell lines

BFP+ K562, BFP+ HER2+ K562, and mCherry+ BT474 cell lines were provided by the lab of Dr. Rogelio Hernandez-Lopez and cultured in RPMI 1640 plus GlutaMAX with 10% FBS. ffLuc+ RFP+ 143B and ffLuc+ GFP+ MG63.3 cell lines were provided by the lab of Dr. Crystal Mackall and cultured in RPMI 1640 plus GlutaMAX with 10% FBS, and 100 U/mL penicillin, and 100mg/mL streptomycin (Gibco #15140122). Lx293T were purchased from Takara Bio USA (#632180) and cultured in DMEM (Life Technologies #10564029) + 10% FBS.

### Arrayed screens

All arrayed screening for SCR CAR T cells were performed in 96-well flat-bottom plates. In the large screen, Seven 96-well plates were created by transforming a pooled combinatorial library into 5-alpha F’ Iq competent *E. coli* (New England Biolabs #C2992H), randomly picking 600 colonies, isolating plasmids by miniprep, and sequencing the variable region of the SCR signaling domains. During plate curation, samples with poor sequencing results were discarded. Samples were run in singlicate with the exception of internal controls. Internal controls included no vector (2 per plate), GFP only (2 per plate), CAR with an SCR containing no motifs (2 per plate), CAR with a non-signaling SCR containing no motifs (2 per plate), SCR containing no motifs (2 per plate), and non-signaling SCR containing no motifs (2 per plate), and a subset of randomly chosen samples which were added to multiple wells during plate curation. In smaller validation arrays all samples were run with three or more replicates.

Prior to addition of cancer cells (approximately 6-10 days after T cell transduction) baseline transduction efficiency and phenotyping (CD45RA, CD62L, CD127, KLRG1, DAPI) was performed via flow cytometry. Samples were challenged (pulsed with cancer cells) every 48 hours for a total of 4 challenges. Upon addition of cancer cells, samples were centrifuged at 100 g for 3 minutes and incubated for 24 hours. At 24 hours, samples were mixed by pipetting and centrifuged at 100 g for 3 minutes. 48 hours after the start of each challenge, samples were mixed by pipetting and 80 μL of 200 μL total was taken for phenotype and killing assessment via flow cytometry. The remaining volume was re-challenged with additional cancer cells in fresh Immunocult-XF media. Within each screen, all samples were pulsed with the same number of new cancer cells at any given challenge. In the large arrayed screen against K562 cell, challenge 1, 2, 3, 4 used a progressive effector to target (E:T) ratio of 1:1, 1:2, 1:4, and 1:8 with 1 being the average T cell number across all samples at the start of the screen, measured by flow cytometry baseline measurement.

Arrayed samples were mixed by pipetting and 80 µL of each sample was transferred from a 96-well flat-bottom plate to a round-bottom plate. Samples were washed in FACS buffer: calcium-free magnesium-free PBS with 3% FBS (Gibco #16140071) and 5mM EDTA (Promega #V4231). A staining solution of antibodies was prepared for staining (for 100 wells) in 3 mL of FACS buffer as follows: 25 µL of CD62L-BV711, 25 µL of CD45RA-PE, 25 µL of CD127-PECy7, and 25 µL of KLRG1-APC. In some validation screens, anti-KLRG1 was excluded and either anti-CD4 or anti-CD8 antibodies were included. Samples were resuspended in 30 µL of staining solution and incubated for 30-60 minutes at room temperature. Cells were then washed twice with FACS buffer. The supernatant was discarded, and the cells were resuspended in 120 µL of FACS buffer containing 1 µg/mL of DAPI. Cells were analyzed by flow cytometry using a ThermoFisher Attune NxT. Antibodies are as follows: APC mouse anti-human KLRG1 clone 13F12F2 (eBioscience #17-9488-42), PE-Cy7 Mouse anti-human IL7Ra clone eBioRDR5 (eBioscience #25-1278-42), BV711 Mouse anti-human CD62L clone DREG56 (eBioscience #407-0629-42), PE Mouse anti-human CD45RA clone HI100 (eBioscience #12-0458-41), APC eFluor780 mouse anti-human CD4 clone RPA-T4 (eBioscience #47-0049-42), and APC mouse anti-human CD8a clone RPA-T8 (eBioscience #17-0088-42).

### Assessment of STAT phosphorylation by flow cytometry

Phosphorylation of STAT1, STAT3, STAT4, and STAT5 were measured in primary human T cells expressing SCRs. STAT phosphorylation was measured for SCRs with different single motifs to assess the contribution of each motif to signaling, different number of alanine residues to assess the effect of box motif orientation on signaling, or different motif combinations to assess the combined effects of motifs on signaling. To quantify SCR-induced STAT phosphorylation, T cells were transferred to cytokine-free Immunocult-XF T Cell Expansion medium and starved of cytokine for 48 hours. Cells were centrifuged and media was removed by flicking. Samples were washed twice with PBS. Cells were then fixed with 1.6% formaldehyde for 15 minutes at room temperature, washed with PBS + 10% FBS, and permeabilized with ice-cold methanol overnight. Cells were washed again with PBS + 10% FBS, then stained with antibodies overnight at 4°C. Stained cells were washed twice with PBS + 10% FBS and resuspended in FACS buffer for flow cytometry data collection. After data collection, channels corresponding to GFP stains were used to select GFP+ T cells for phosphoSTAT analysis. The following antibodies were used to detect intracellular proteins: PE-anti-phospho-STAT1 (Tyr701) (clone KIKSI0803, eBioscience #12-9008-42), Pacific Blue-anti-phospho-STAT3 (Tyr705) (clone 4/P-STAT3, BD Biosciences #560312), PE-CF594-anti-phospho-STAT4 (Tyr693) (clone 38/p-stat4, BD Biosciences #567631), R718-anti-phospho-STAT5 (Tyr694) (clone 47/Stat5, BD Biosciences #566978), PE-anti-GFP (clone FM264G, BioLegend #338003), and eFluor660-anti-GFP (clone 5F12.4, eBioscience #50-6498-82).

### Mice

All mouse experimental procedures were conducted according to Institutional Animal Care and Use Committee (IACUC)–approved protocols (AN183960-02A). The experiments were planned and independently performed by Stanford University based on Stanford University studies. Female immunocompromised NOD-SCID-*Il2rg*^-/-^ (NSG) mice were obtained from The Jackson Laboratory. At time of tumor injections, mice were approximately 8 weeks old. Mice were humanely euthanized when an IACUC-approved endpoint (hunching, neurological impairments such as circling, ataxia, paralysis, limping, head tilt, balance problems, seizures, tumor diameter greater than 17 mm) was reached (5 mice per group).

#### *143B* Osteosarcoma

On Day 1, mice were injected subcutaneously with 0.7 × 10^6^ 143B osteosarcoma cells. On Day 5, mice were injected with 5×10^6^ T cells via tail vein injection. Osteosarcoma progression was measured by caliper twice per week.

### Machine learning data preparation

The array data include three types of independent variables: plate number (to account for batch effects), GFP+ T cell count and percent of T cells that are GFP+ (to account for T cell dose), and a vector describing the SCR motif combination (signaling domain composition). Before training, the input data were one-hot encoded. Each motif position was described by a vector of fourteen 0s, and one 0 in each vector was replaced with a 1 corresponding to the absence of a motif (replace the first 0 with 1), the presence of a motif (replace the 0 equal to the part number + 1 with 1; S1 is represented by a 1 in the second position, S12 is represented by a 1 in the thirteenth position, and SL is represented by a 1 in the 14^th^ position). During synthesis of the 4-motif SCR library, several SCRs containing 5 or 6 motifs were created. Rather than discard the data for these SCRs, we allowed up to 6 motif positions in the model, for a total of 6 vectors. Values for GFP+ T cell count, and percent of T cells that are GFP+ were normalized to vary from 0 to 1 with 0 corresponding to the minimum value observed in the training data and 1 corresponding to the maximum value observed in the training data. Data were randomly split into a training set containing 489 examples and a test set containing 50 examples.

### Machine learning model training

In this work, we used a Convolutional Neural Networks (CNN), followed by a Long Short-Term Memory (LSTM) network together with fully connected layers. The code is implemented in *Mathematica* (Wolfram). The Mathematica analysis is described below. The neural network uses plate number (for batch effects), GFP+ T cell count and percent of T cells that are GFP+ (T cell dose), and a vector describing the SCR motif combination (signaling domain composition) as inputs and outputs one value corresponding to one of the phenotypes (cytotoxicity, memory, or unstimulated proliferation).

The plate number is first passed through an embedding layer which outputs a vector of length 14 (to match the length of the vectors describing the motifs) which is then passed to a fully connected layer with ReLU activation and then to the catenation layer. The two numbers describing T cell dose are first passed through a fully connected layer with linear activation which outputs a vector of length 14 (to match the length of the vectors describing the motifs), which is then passed to a fully connected layer with ReLU activation, then to the catenation layer. The motif vectors are first passed through a fully connected layer with linear activation then to a fully connected layer with ReLU activation and to a catenation layer.

Between the catenation layer and output, there is 1 convolutional layer, 1 LSTM layer, 1 dropout layer, and 3 fully connected layers with linear activation, and 1 fully connected layer with ReLU activation. We used dropout regularization to prevent over-fitting. For training in Mathematica, we used mean squared error loss and ADAM optimization algorithm with automatic learning rate, and training over 200 iterations.

### Selection of neural network hyperparameters

We tuned the hyperparameters for layers in the neural networks to find optimal hyperparameters for the cytotoxicity, memory, and unstimulated proliferation datasets. The tuned hyperparameters include convolutional layers filters (20, 50), kernel size (3, 5); LSTM layer units (6, 12, 18), and units for two fully connected layers ((16, 64, 128) and (1, 4, 16)).

Hyperparameters were tuned as follows: We performed a grid search of hyperparameters and scored each parameter set by 10-fold cross validation of the training set. The best-performing 10 hyperparameter sets for each dataset (cytotoxicity, memory, and unstimulated proliferation) were selected using the K-fold averaging cross validation (ACV) method and used to train 10 neural networks whose outputs were then averaged (*50*). Due to the stochastic nature of network initialization and dropout, as well as the availability of a limited training set, every neural network is unique in terms of the parameterization of the network connections (*51, 52*). To obtain consensus predictions we averaged the predictions from the 10 neural networks trained on each phenotype.

Hyperparameters for final neural networks are available in supplementary tables.

### Simulation of SCR libraries

The trained neural networks were used to simulate the cytotoxicity, memory, and unstimulated proliferation for the 30,940 combinations of 1, 2, 3, or 4 variable motifs at a fixed initial GFP + T cell count of 5869 at 67 %GFP+ with batch label set to mimic Plate 1.

### Motif enrichment analysis

Motif enrichment analysis was performed in *Mathematica* (Wolfram) to assess contribution of each signaling motif (variable part) to SCR CAR T cell cytotoxicity and memory. For each of 10 models, 30,940 SCRs were sorted by predicted cytotoxicity. The count of each motif or motif pair in the top 50% (most cytotoxic) was divided by the count in the bottom 50% (least cytotoxic) SCRs to calculate the fold change (FC). The Log2FC was calculated from the FC. Each p-value was calculated by performing a Student’s t-test to compare the observed FC values from 10 model predictions to the 10 expected FC values for the null hypothesis (null expected FC = 1). The - Log10p-values were calculated from the p-values.

### In silico mapping of signaling landscape

Derivations can be found in Appendix I. To predict the percent of cells that are pSTAT1, pSTAT3, pSTAT4, and pSTAT5 positive when transduced with a given SCR, we assume that the percent of positive cells is equal to 100*(K_p_/(1+K_p_)). For an SCR with no motifs (SCR0), K_p_ = K_i_, the intrinsic phosphorylation constant. We assume there is a unique Ki for each pSTAT. For SCRs with a single motif, K_p_ = K_i_*K_m_ where Km is the motif-specific phosphorylation constant. If a motif increases phosphorylation of a pSTAT, Km is greater than 1. If a motif reduces phosphorylation of a pSTAT, Km is less than 1. For an SCR with multiple motifs, K_p_ = K_i_*Product(K_m*j*_) where K_m*j*_ are the K_m_ values for each motif in the SCR. We do not consider position or any additional interactions between motifs.

To obtain K_i_ and K_m_ values, we fit the model described in Fig. 5D. to experimental measurements of phosphorylation for SCRs with single motifs (Fig. 1E) and SCRs with four motifs (fig. S5A). During fitting, the floating parameters were ΔGi values for the K_i_ corresponding to pSTAT1, pSTAT3, pSTAT4, and pSTAT5. We fit for ΔGi values rather than K_i_ to avoid unrealistic negative K_i_ values. Here, ΔG_i_=-RTln(K_i_) where R = 8.314 J mol^-1^ K^-1^ and T = 310K. During the fit Km values were calculated using the equation K_m_ = -x/((−100+x)K_i_) where x is the measured percent of T cells positive for pSTAT. This is a rearrangement of the equation x = 100*Ki* K_m_/(1+Ki* K_m_) in Figure 5D. Fitted Ki and calculated K_m_ values were used to predict pSTAT values for SCRs with four motifs. The object of the fit minimized the sum of squared errors between the measured and predicted pSTAT values for SCRs with four motifs (Fig. 5E).

To map signaling along the Pareto front, an arc was swept from 0.5 Pi to Pi in steps of 0.1 Pi and SCRs with the greatest cytotoxicity (least surviving K562s) in each step (up to 20 SCRs per step) were labeled as being on the Pareto front. For the 476 SCRs along the Pareto front, we used th fitted model to predict the percent pSTAT1, pSTAT3, and pSTAT5 positive cells as described above. For the motif counts in Figure 5G We totaled the number of motifs in each SCR expected to bind PI3K, Grb2, Shc, and PLCγ1 based on the presence of consensus motifs shown in Figure 1F.

Visualizations of signaling along SCR trajectories (Fig. 4F through G and Fig. 5E through F) represent smoothed predictions generated as follows: First, we calculated the running average of predicted pSTAT signaling (or motif count) in runs of 15 SCRs. Second, we selected 7 SCRs spaced SCRs evenly along the SCR trajectory. Next, the mean and standard error of pSTAT signaling (or motif count) for the 50 SCRs nearest to the 7 SCRs. Bands were plotted in *Mathematica 14.1.0.0* using ListLinePlot with InterpolationOrder = 2. The center of the band represents the mean, and the upper and lower band represent the mean plus or minus the standard error. An overlay of moving averages and bands can be found in figure S5E-H. We made similar predictions of pSTAT signaling for SCRs along the center diagonal (fig. S5I) and SCRs far from the Pareto front (fig. S5J).

Predicted maps of pSTAT signaling for SCRs in the predicted library (fig. S5K) were created in *Mathematica 14.1.0.0* as follows. We rescaled the predicted library cytotoxicity and memory values to range from 0 to 1. We then created a grid of 51 x 51 waypoints ranging from 0 to 1 in steps of 0.02. For each waypoint, we calculated the z value as the average predicted pSTAT1, 3, 4, or 5 signaling for the 50 nearest (and within a radius of 0.05) rescaled predicted library datapoints. Waypoints with no predicted SCRs within a radius of 0.05 were discarded. For the resulting four (one for each pSTAT) grids of 2601 waypoints, the x and y values were then rescaled to match the cytotoxicity and memory values of the predicted library. The color of each waypoint represents the z value corresponding to average predicted pSTAT signaling of nearby SCRs.

---

**Figure S1.**
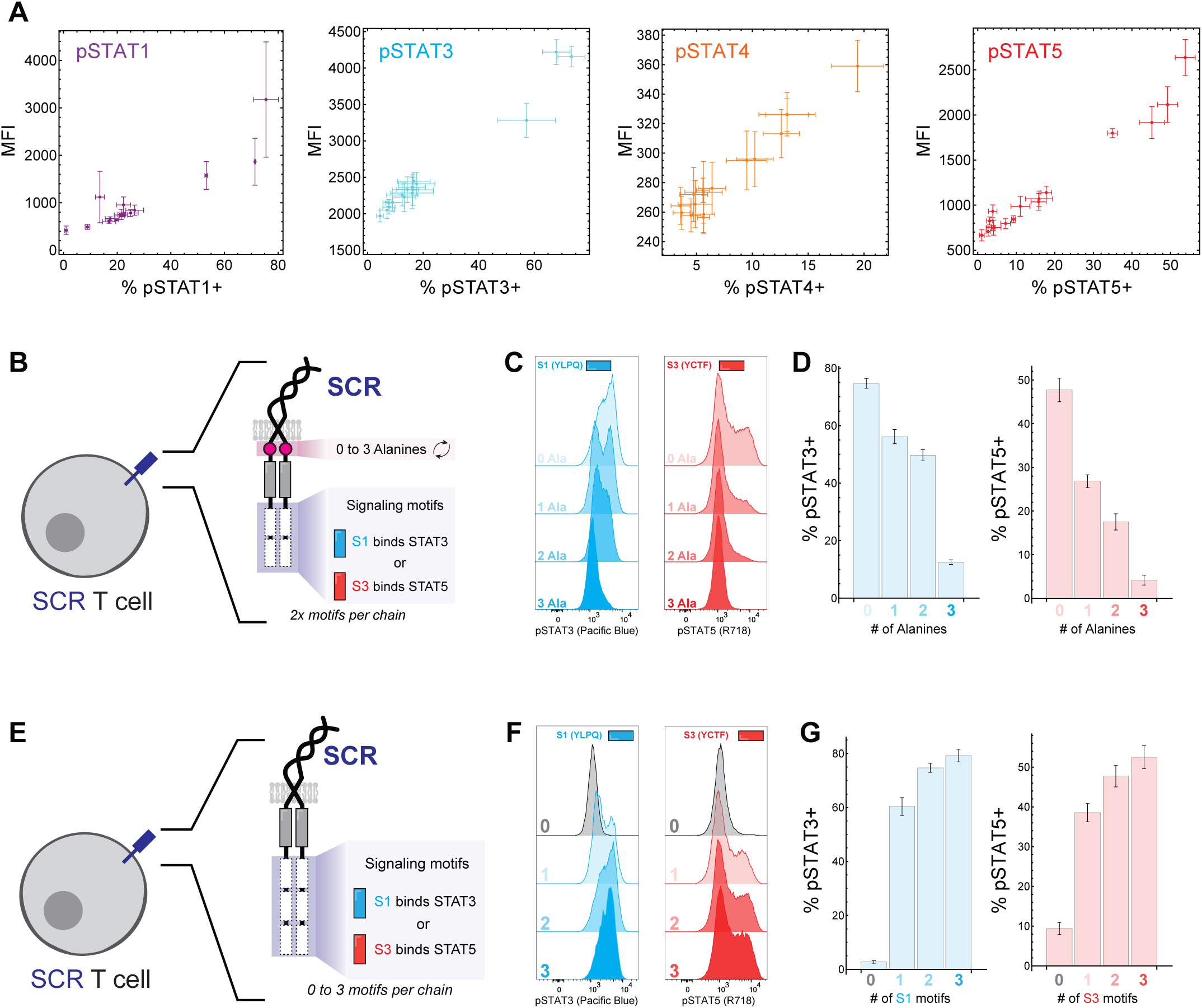
Constitutively active synthetic cytokine receptors (SCR) induce signaling motif-dependent signaling and T cell phenotype. A) The percent of pSTAT positive cells and the mean fluorescence intensity (MFI) upon pSTAT staining are proportional. Flow cytometry was used to measure pSTAT1, 3, 4, and 5 in primary human T cells transduced with SCRs containing a single motif or no motif. B) Primary human T cells were engineered to express a homodimeric SCR either two copies of motif S1 or two copies of motif S3. SCRs contained 0, 1, 2, or 3 alanine residues inserted between the transmembrane domain and signaling domain. C) Flow cytometry was used to measure phosphorylation of STAT3 in T cells with SCRs containing motif S1, or phosphorylation of STAT5 in T cells with SCRs containing motif S3. D) Quantification of the percent of T cells positive for pSTAT3 or pSTAT5 as function of alanine residue number. E) Primary human T cells were engineered to express a homodimeric SCR 0, 1, 2, or 3 copies of motif S1 or motif S3. F) Flow cytometry was used to measure phosphorylation of STAT3 in T cells with SCRs containing motif S1, or phosphorylation of STAT5 in T cells with SCRs containing motif S3. G) Quantification of the percent of T cells positive for pSTAT3 or pSTAT5 as function of motif copy number. Values and error bars in A, C, and F represent mean and standard error of 3 replicates.

**Figure S2.**
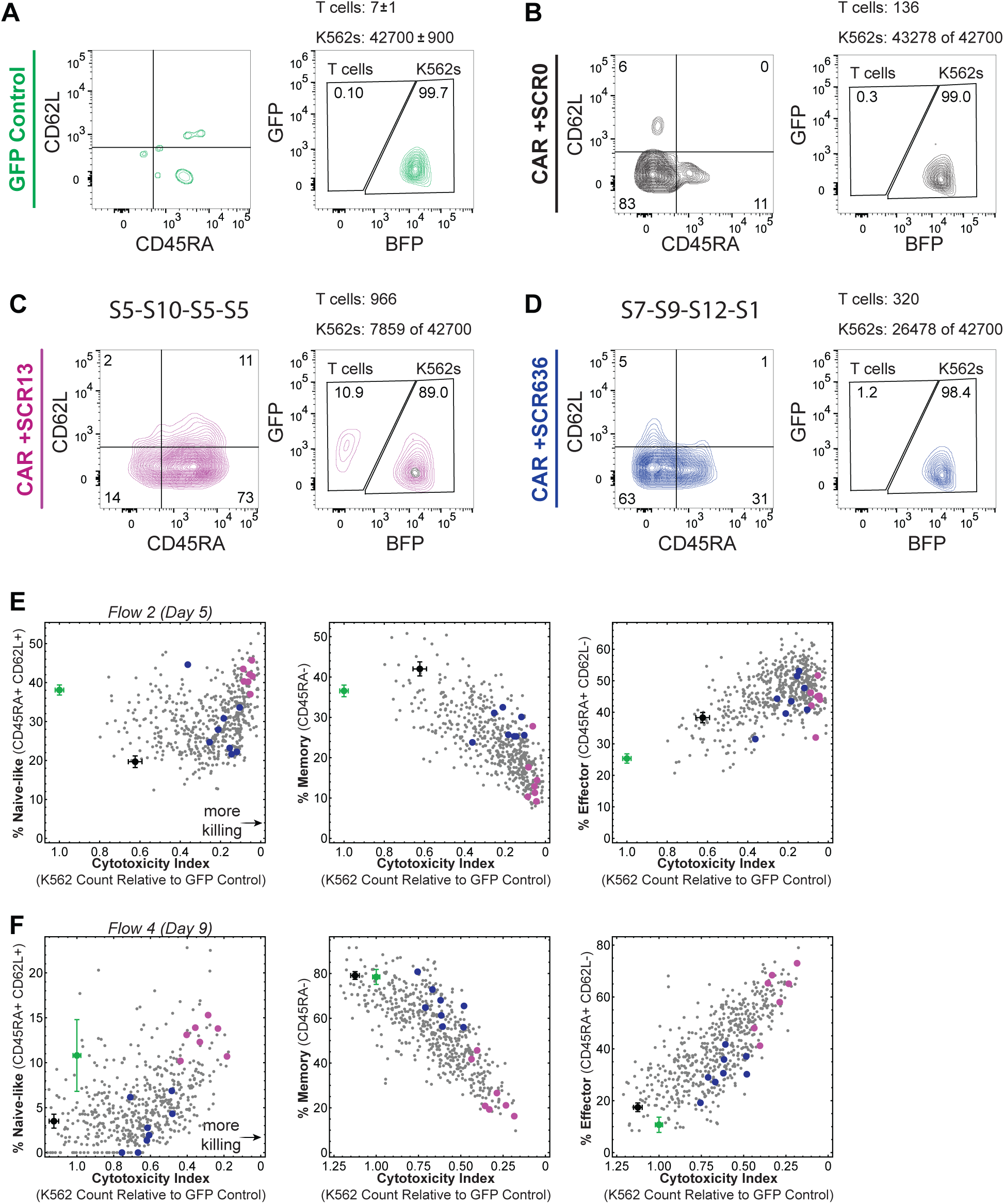
An SCR library with synthetic combinations of signaling motifs generates diverse CAR T cell states. A-D) Representative flow cytometry plots of T cells expressing GFP control (A), SCR0 + CAR (B), SCR13 + CAR (C), or SCR636 + CAR in co-culture with HER2+ K562 target cells. Flow cytometry data was collected on day 9 of the co-culture. E) Cytotoxicity of SCR CAR T cells and controls versus naïve-like, memory, or effector T cell population on day 5 of the co-culture with HER2+ K562 cells. F) Cytotoxicity of SCR CAR T cells and controls versus naïve-like, memory, or effector T cell population on day 9 of the co-culture with HER2+ K562 cells.

**Figure S3.**
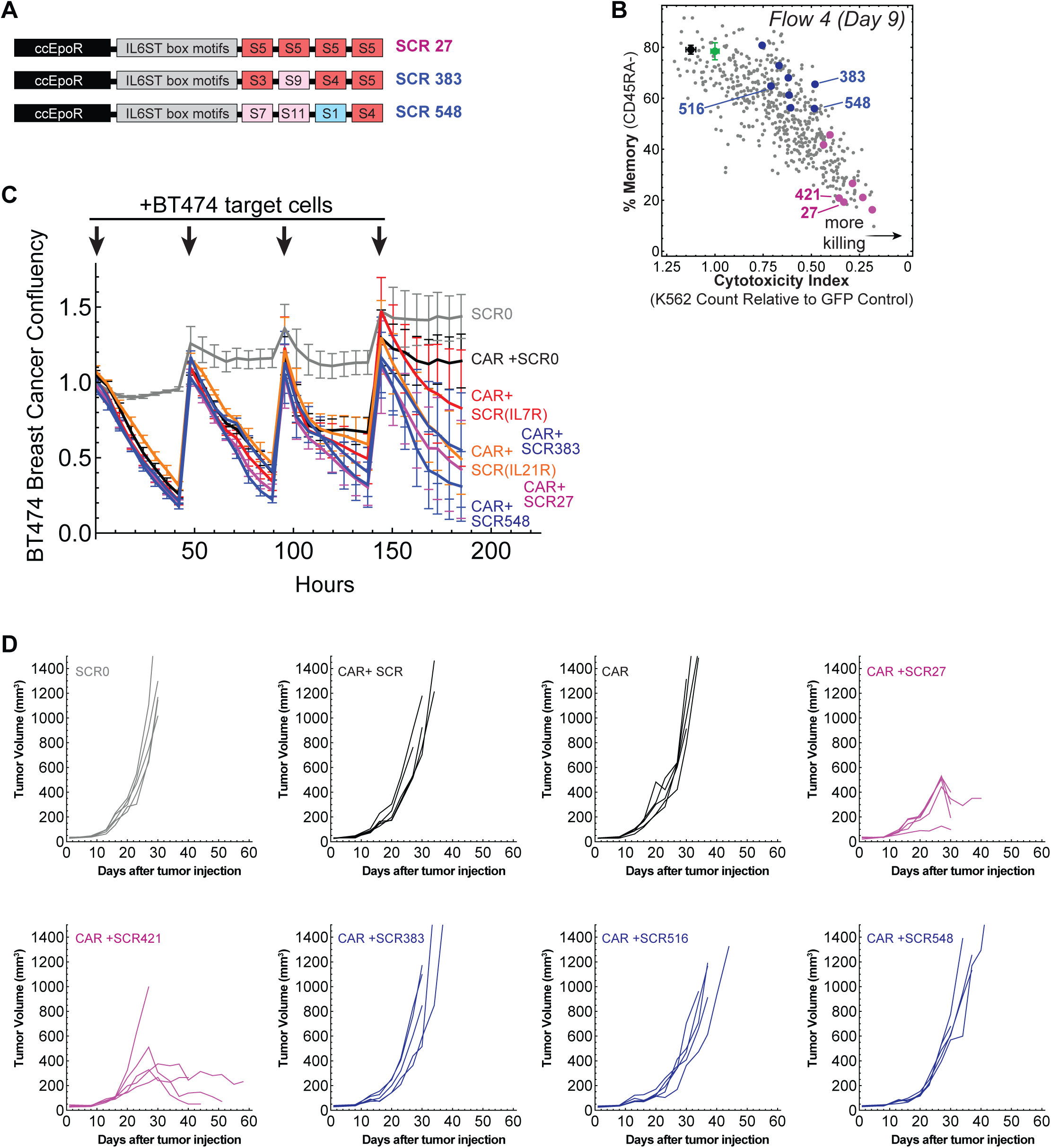
SCRs with synthetic signaling domains improve CAR T cell anti-tumor activity. A) Schematics of SCR27, SCR383, and SCR548. B) Day 9 flow cytometry measurements of cytotoxicity and memory for the library data in Figure 2. SCRs of interest (27, 383, 421, 516, and 548) are labeled. C) SCR CAR T cells were pulsed with mCherry+ BT474 breast cancer cells at 0, 48, 96, and 144 hours. Confluency of BT474 cells was monitored by live cell imaging to assess SCR CAR T cell cytotoxicity. D) NSG mice were injected with 0.7×10^6^ 143B on day 1 and 5×10^6^ T cells on day 5. Tumor volumes were measured two to three times weekly by caliper. n = 5 mice per group. Values and error bars in B represent mean and standard error of 3 replicates.

**Figure S4.**
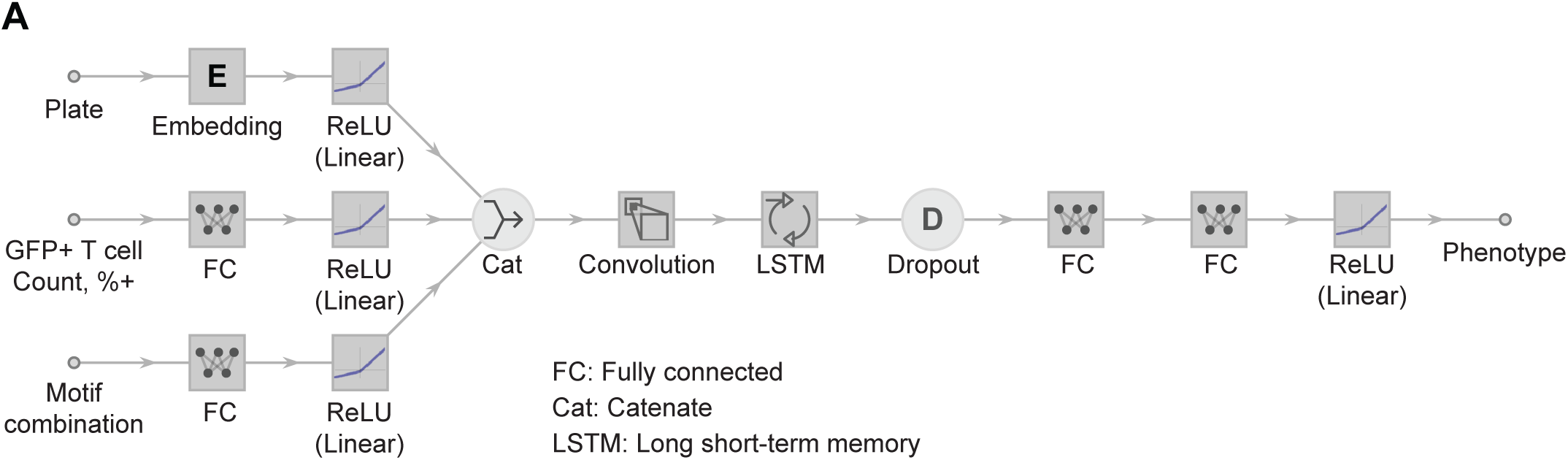
Neural networks predict CAR T cell memory, cytotoxicity, and autonomous proliferation encoded by SCR signaling motif combinations. A) Detailed schematic of the neural network used to predict SCR CAR T cell phenotype.

**Figure S5.**
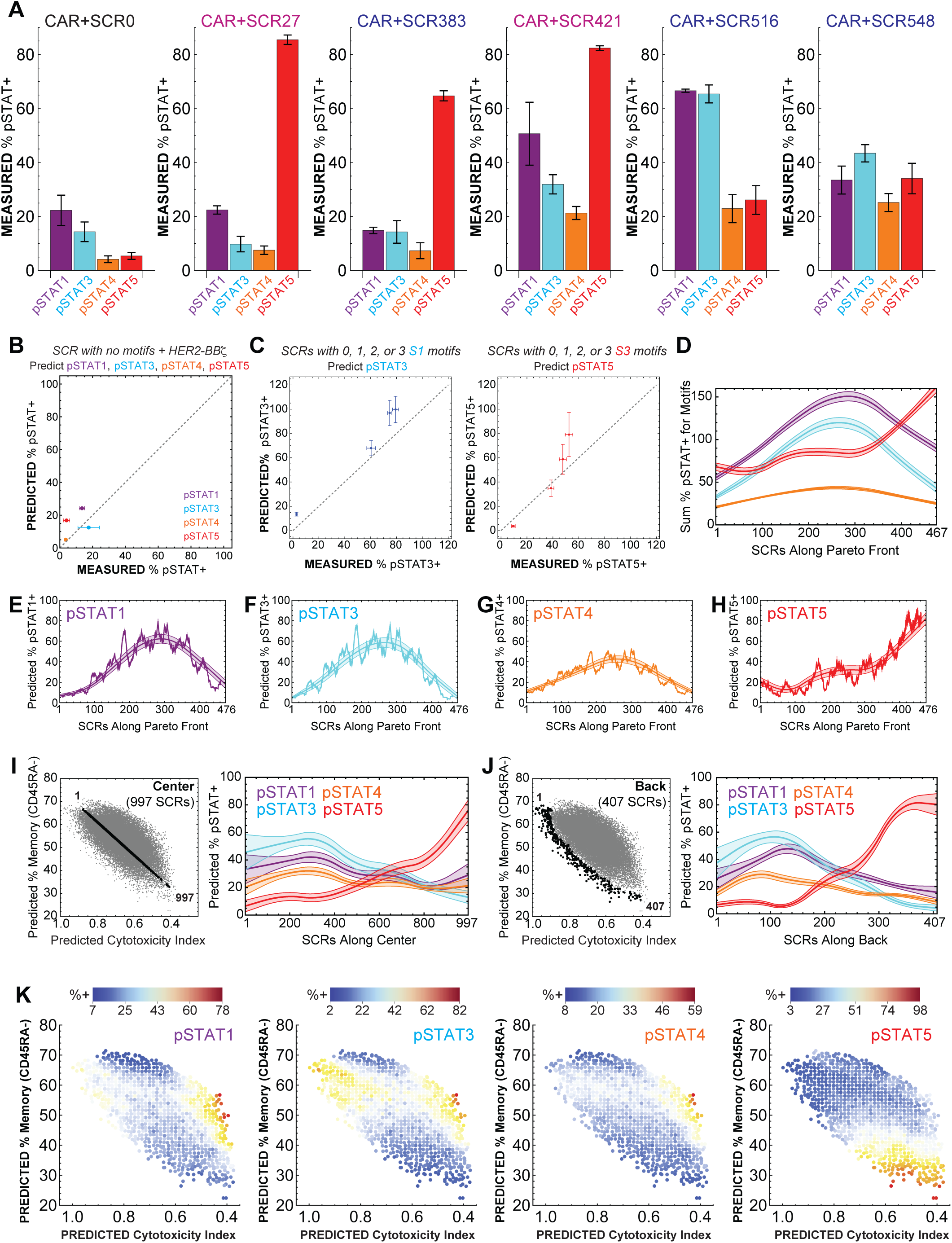
A continuous landscape of JAK/STAT signaling encodes a corresponding landscape of CAR T cell phenotype. A) Quantification of the percent of SCR CAR T cells positive for pSTAT1, 3, 4, or 5. Mean and standard error of n=3 measurements. B) Measured and predicted values for pSTAT signaling for SCRs with no signaling motifs. Predicted pSTAT signaling was calculated from fitted K*i* values. C) Measured and predicted values for pSTAT signaling for SCRs with 0, 1, 2, or 3 copies of signaling motifs S1 or S3. Predicted pSTAT signaling was calculated from fitted K*i* values and K*m* values. D) Sum of pSTAT signaling for motifs in SCRs along the Pareto front. The measured pSTAT signaling for each motif (Fig. 1E) was used to calculate the sum of pSTAT signaling. E-H) The fitted model in Fig. 4D was used to predict pSTAT1, 3, 4, and 5 signaling in SCR CAR T cells along the Pareto front. Jagged lines represent the running average and bands representing mean plus or minus standard error. I-J) The fitted model of SCR-induced pSTAT signaling was used to predict the pSTAT1, 3, 4, and 5 signaling for SCRs along the center diagonal (I) or far from the Pareto front (J). K) The fitted model of SCR-induced pSTAT signaling was used to predict the pSTAT1, 3, 4, and 5 signaling for SCRs in the predicted library. The color of each point represents the average predicted pSTAT signaling of nearby SCRs.

**Table S1.** Hyperparameters for neural networks trained on day 9 cytotoxicity data. Hyperparameters were selected by grid search as described in the methods.

| Neural Network Model | Convolutional Layer Filters | Convolutional Layer Kernel Size | LSTM Layer Units | Fully Connected Layer Units | Fully Connected Layer Units |
| --- | --- | --- | --- | --- | --- |
| NN1: | 50 | 5 | 6 | 16 | 16 |
| NN2: | 50 | 5 | 6 | 16 | 1 |
| NN3: | 20 | 5 | 18 | 64 | 4 |
| NN4: | 20 | 5 | 18 | 64 | 1 |
| NN5: | 50 | 5 | 18 | 128 | 4 |
| NN6: | 20 | 3 | 12 | 64 | 16 |
| NN7: | 50 | 3 | 12 | 128 | 1 |
| NN8: | 50 | 3 | 18 | 16 | 4 |
| NN9: | 20 | 5 | 18 | 16 | 4 |
| NN10: | 50 | 3 | 12 | 64 | 1 |

**Table S2.**
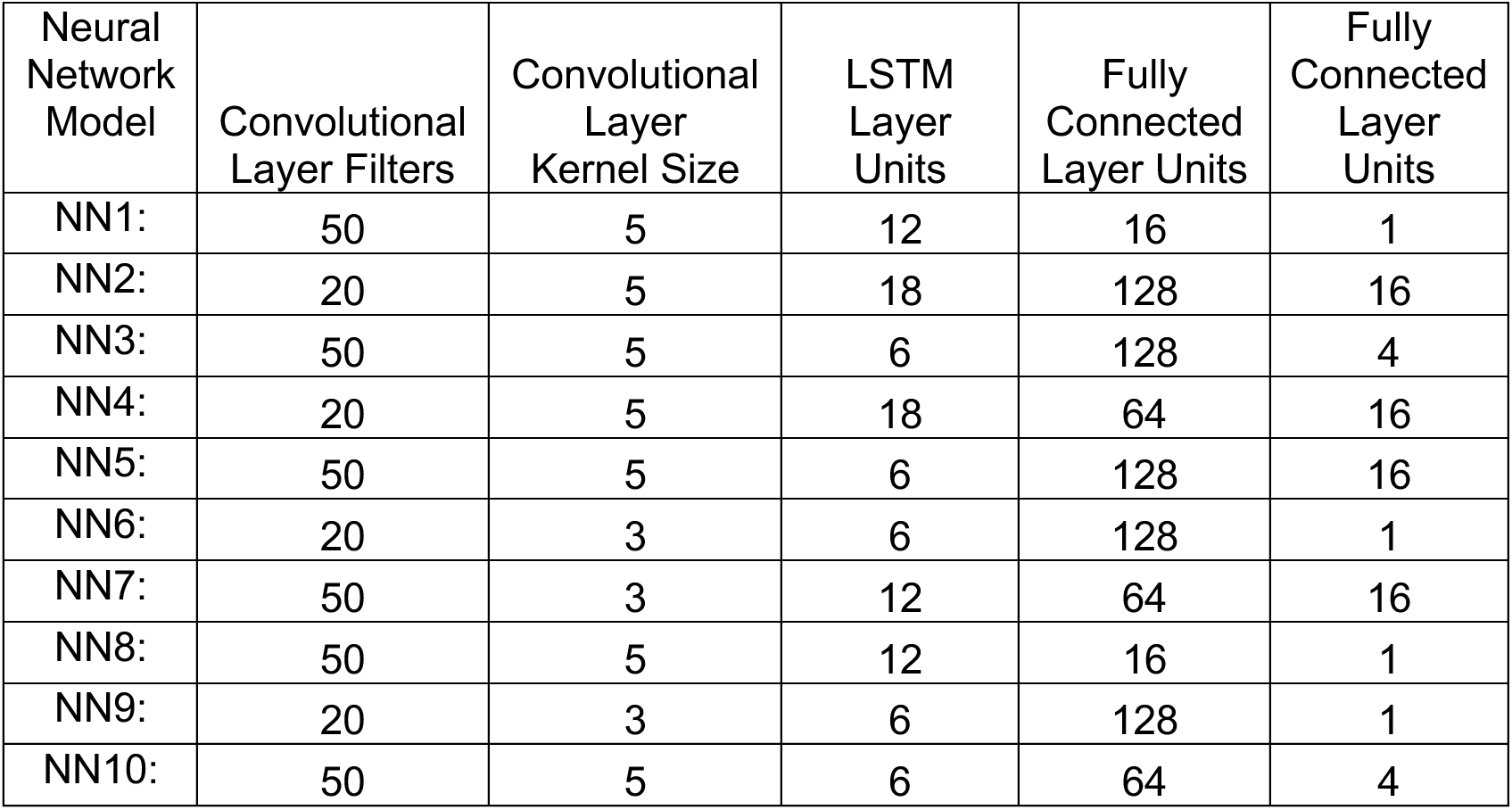
Hyperparameters for neural networks trained on day 9 memory data. Hyperparameters were selected by grid search as described in the methods.

**Table S3.** Hyperparameters for neural networks trained on unstimulated proliferation data. Hyperparameters were selected by grid search as described in the methods.

| Neural Network Model | Convolutional Layer Filters | Convolutional Layer Kernel Size | LSTM Layer Units | Fully Connected Layer Units | Fully Connected Layer Units |
| --- | --- | --- | --- | --- | --- |
| NN1: | 20 | 3 | 18 | 16 | 1 |
| NN2: | 20 | 3 | 12 | 128 | 4 |
| NN3: | 20 | 5 | 18 | 128 | 4 |
| NN4: | 50 | 5 | 6 | 64 | 1 |
| NN5: | 20 | 5 | 6 | 128 | 4 |
| NN6: | 20 | 5 | 12 | 128 | 1 |
| NN7: | 20 | 5 | 6 | 64 | 1 |
| NN8: | 50 | 5 | 12 | 16 | 1 |
| NN9: | 20 | 3 | 18 | 128 | 1 |
| NN10: | 50 | 5 | 18 | 64 | 4 |

Antibody Table.

Antibody Table
| Target | Clone | Source Animal | Fluorophore | Manufacturer | Vendor | Product Number |
| --- | --- | --- | --- | --- | --- | --- |
| CD45RA (human) | HI100 | Mouse | PE | eBioscience | Invitrogen | 12-0458-41 |
| CD62L (human) | DREG56 | Mouse | Brilliant Violet 711 | eBioscience | Invitrogen | 407-0629-42 |
| IL7Ra (human) | eBioRDR5 | Mouse | PE-Cy7 | eBioscience | Invitrogen | 25-1278-42 |
| KLRG1 (human) | 13F12F2 | Mouse | APC | eBioscience | Invitrogen | 17-9488-42 |
| pSTAT1, Y701 (human) | KIKSI0803 | Mouse | PE | eBioscience | Invitrogen | 12-9008-42 |
| pSTAT3, Y705 (human) | 4/P-STAT3 | Mouse | Pacific Blue | BD Bioscience | Fisher Scientific | 560312 |
| pSTAT5, Y694 (human) | 47/Stat5 | Mouse | R718 | BD Bioscience | Fisher Scientific | 566978 |
| pAkt (human) | M89-61 | Mouse | Alexa Fluor 647 | BD Bioscience | Fisher Scientific | 560343 |
| pNFkB (human) | B33B4WP | Mouse | PE-Cy7 | eBioscience | Invitrogen | 25-9863-42 |
| pErk (human) | 6B8B69 | Mouse | Brilliant Violet 421 | BioLegend | BioLegend | 369509 |
| myc |  |  |  |  |  |  |
| pSTAT4, Y693 (human) | 38/p-Stat4 | Mouse | PE-CF594 | BD Bioscience | Fisher Scientific | 567631 |
| GFP | FM264G | Rat | PE | BioLegend | BioLegend | 338003 |
| GFP | 5F12.4 | Rat | eFluor 660 | eBioscience | Invitrogen | 50-6498-82 |
| CD4 | RPA-T4 | mouse | APC eFluor780 | eBioscience | Invitrogen | 47-0049-42 |
| CD8a | RPA-T8 | mouse | APC | eBioscience | Invitrogen | 17-0088-41 |

## Data Availability

Datasets, neural network predictions, and Mathematica notebooks will be available upon publication of peer-reviewed article. Plasmids encoding the SCR and signaling motifs be available from Addgene.

## Acknowledgments

We thank Adam Stevens and Anusha Kalbasi for illuminating feedback. We thank the Bassik, Barsh, Baker, Kirkegaard, Sherlock, and Hernandez-Lopez labs for generous sharing of equipment and reagents. This work was supported by grants from The Hypothesis Fund, The Shurl and Kay Curci Foundation, the Parker Institute for Cancer Immunotherapy, Stanford C-ShaRP, the Stanford Cancer Institute, and the Stanford Genetics Department. AS and LS are supported by the NSF Graduate Research Fellowship Program.

## Author Contributions

WC, ANB, and KGD conceptualized the study. WC and KGD wrote and edited the manuscript with input from all authors. WC, JL, ANB, JCL, PX, KH, EEC, AS, LS, KO, and NP performed experiments. WC, JL, and KGD set up computational pipelines and analyzed the data. WC, ANB, ES, and KGD guided experiments and data analysis.

